# A Dual Role for the PP2A Phosphatase in Hippo Signalling Regulation

**DOI:** 10.1101/2024.11.14.623552

**Authors:** Aashika Sekar, Alberto Rizzo, Elodie Sins, Alexander D. Fulford, Paulo S. Ribeiro

## Abstract

Hippo signalling is an evolutionarily conserved pathway that regulates tissue growth. The FERM domain protein Expanded plays a crucial role in integrating polarity cues to activate the Hippo pathway. Previous work has shown that the apicobasal polarity protein Crumbs can limit Hippo activity by promoting the phosphorylation and degradation of Expanded. Here, we provide evidence that PP2A^Wrd^ can counteract the effects of Crumbs, by dephosphorylating and stabilising Expanded. Indeed, we demonstrate that the PP2A^Wrd^ holoenzyme can increase Hippo signalling activity, in contrast to the previously established Hippo pathway inhibitory role of the PP2A^Cka^-containing STRIPAK complex. We also uncover a role for PP2A^Wrd^ and PP2A^Tws^ in the regulation of Expanded proteostasis. Remarkably, the upstream Hippo regulator, Kibra interacts with PP2A^Wrd^ and prevents Expanded degradation. However, Kibra is unable to antagonise Crumbs-mediated Expanded regulation, in agreement with the previously established role of Crumbs in inhibiting Kibra function. Overall, our work characterises a novel Hippo-activating role for PP2A in the stabilisation of Expanded and provides new insights into how PP2A tightly controls Hippo activity in response to polarity stimuli.

## Introduction

In multicellular organisms, achieving the appropriate final organ size and development of proportionate adult organisms require precise orchestration of cell growth, division and death, that can be fine-tuned by intrinsic and extrinsic cues (Yu, Zhao and Guan, 2015). The evolutionarily conserved Hippo (Hpo) pathway is a such key tissue growth regulator and plays a role in maintaining tissue architecture especially in epithelial tissues (Genevet and Tapon, 2011; Schroeder and Halder, 2012; Karaman and Halder, 2018). At its core, the Hippo pathway is comprised of a kinase cascade that includes the *Drosophila* kinases Hpo and Warts (Wts) and their corresponding adaptor proteins, Salvador (Sav) and Mob as tumour suppressor (Mats) (Fulford, Tapon and Ribeiro, 2018; Ma *et al*., 2019; Zheng and Pan, 2019; Fu *et al*., 2022). Activation of the Hpo kinase cascade culminates in the phosphorylation and subsequent cytoplasmic sequestration of the pro-growth transcriptional co-activator Yorkie (Yki), resulting in growth repression (Huang *et al*., 2005; Zhao *et al*., 2007). When the pathway is inactive, Yki translocates to the nucleus and interacts with its cognate transcription factor Scalloped to promote expression of genes that broadly promote cell proliferation and inhibit apoptosis, such as *cyclin E and diap1* (Wu *et al*., 2008; Zhang *et al*., 2008). To prevent unrestricted Yki-mediated growth and maintain tissue homeostasis, a negative feedback loop is incorporated into the Hpo pathway, whereby the expression of upstream activators such as *expanded* (*ex*), *kibra* (*kib*) and *merlin* (*mer*) is controlled by Yki itself, thereby limiting Yki activity (Hamaratoglu *et al*., 2006; Genevet *et al*., 2010; Park and Hansen, 2021). Importantly, deregulation of Hpo pathway components is associated with disease development and progression, including multiple cancer types (Yu, Zhao and Guan, 2015; Fu *et al*., 2022).

Unlike conventional signalling pathways that depend on specific ligand/receptor complexes, Hpo signalling senses and responds to diverse upstream local and external signals that impact on epithelial cell structure and function (Fulford, Tapon and Ribeiro, 2018; Fu *et al*., 2022). One key regulator is the Striatin-interacting phosphatase and kinase (STRIPAK) complex (Glatter *et al*., 2009; Goudreault *et al*., 2009; Ribeiro *et al*., 2010; Liu *et al*., 2016; Bae *et al*., 2017; Zheng *et al*., 2017; Chen *et al*., 2019), a large, multicomponent protein complex that includes the serine/threonine phosphatase, protein phosphatase 2A (PP2A) (Jeong *et al*., 2021). PP2A is a trimeric holoenzyme comprising the catalytic subunit Microtubule star (Mts), the adaptor subunit PP2A-29B, and a variable regulatory B subunit that specifies substrate recruitment. The *Drosophila* genome encodes six PP2A regulatory subunits, which are classified into four different families: B (Twins (Tws)), B’ (Well-rounded (Wrd), Widerborst (Wdb) and CG32568), B’’ (CG4733) and B’’’ (Connector of kinase to AP-1 (Cka)). Cka, the *Drosophila* orthologue of mammalian Striatins, is the PP2A regulatory subunit present in the STRIPAK complex (Jeong *et al*., 2021). SLMAP, a component of the STRIPAK complex, recruits Hpo and promotes dephosphorylation of its activation loop, counteracting Hpo autophosphorylation and Tao-1 kinase-mediated Hpo phosphorylation, thereby inhibiting the activity of the Hpo kinase cascade (Genevet *et al*., 2010; Ribeiro *et al*., 2010; Boggiano, Vanderzalm and Fehon, 2011; Zheng *et al*., 2017). RASSF, a known Hpo pathway antagonist that competes with Sav for Hpo binding (Polesello *et al*., 2006), has been shown to associate with STRIPAK and mediate its interaction with Hpo (Ribeiro *et al*., 2010). It has also been reported that SAV1 (Sav mammalian orthologue) can inhibit STRIPAK-mediated dephosphorylation of MST2 (Hpo mammalian orthologue) by directly binding to components of the STRIPAK complex, including SLMAP (Bae *et al*., 2017).

STRIPAK has also been shown to relay hormonal and innate immunity cues to the Hpo pathway and is predicted to have an oncogenic role, since increased STRIPAK activity resulted in enhanced Yki/YAP function (Shi, Jiao and Zhou, 2016; Chen *et al*., 2019; Yang *et al*., 2024).

Another important Hpo pathway upstream regulator is the FERM domain-containing protein Ex, which integrates apicobasal polarity cues to regulate tissue architecture and growth via the Hpo pathway (Genevet and Tapon, 2011; Fulford, Tapon and Ribeiro, 2018). Ex can promote Tao-1-mediated Hpo phosphorylation by regulating the localisation of Schwannomin interacting protein 1 (Schip1), which is essential for the recruitment of the Tao-1 kinase to Hpo (Boggiano, Vanderzalm and Fehon, 2011; Poon *et al*., 2011; Chung, Augustine and Choi, 2016). Ex also acts as a scaffold to recruit Wts to Hpo, promoting Wts phosphorylation (Sun, Reddy and Irvine, 2015). Additionally, the PPxY motifs of Ex can directly interact with the WW domains of Yki to tether Yki to the apical membrane and inhibit its activity in a Hpo-independent manner (Badouel *et al*., 2009; Oh, Reddy and Irvine, 2009). The transmembrane apicobasal polarity determinant Crumbs (Crb) recruits Ex to the apical membrane by directly interacting with its FERM domain, thereby facilitating Hpo pathway activation by Ex (Chen *et al*., 2010; Grzeschik *et al*., 2010; Ling *et al*., 2010; Robinson *et al*., 2010; Hafezi, Bosch and Hariharan, 2012). Apically-localised Ex interacts with the upstream Hpo regulators, Kibra (Kib) and Merlin (Mer) to regulate the Hpo pathway (McCartney *et al*., 2000; Baumgartner *et al*., 2010; Genevet *et al*., 2010; Yu *et al*., 2010). Interestingly, Crb also limits Ex activity by modulating Ex protein turnover. Crb promotes phosphorylation of the N-terminus of Ex by Casein Kinase 1 (CKI) family kinases (Fulford *et al*., 2019), which then subsequently results in Ex ubiquitylation and degradation by the SCF^Slimb/β-TrCP^ E3 ubiquitin ligase complex (Slmb) (Ribeiro *et al*., 2014). Slmb also promotes Ex degradation by interacting directly with its C-terminus (Zhang *et al*., 2015). In parallel, Crb mediates the interaction between the E3 ubiquitin ligase Plenty of SH3s (POSH) and the C-terminus region of Ex resulting in ubiquitylation and subsequent degradation of Ex (Ma *et al*., 2018). The E2 ubiquitin ligase, Bruce, has been shown to act synergistically with POSH to promote Ex degradation (Song and Ma, 2023). Thus, the Crb/Ex axis is a crucial regulatory node linking epithelial polarity inputs and Hpo signalling. However, whether Crb-mediated phosphorylation and degradation of Ex can be antagonised to activate and/or maintain Hpo pathway activity is currently unknown. Identifying this mechanism is vital for understanding how tissue homeostasis is maintained by the Crb/Ex nexus of the Hpo pathway.

Here, we demonstrate that PP2A has a dual role in Hpo signalling regulation. In addition to its previously described inhibitory role as part of the STRIPAK complex, PP2A can act as a Hpo pathway activator. PP2A^Wrd^ (PP2A holoenzyme with the Wrd regulatory subunit) dephosphorylates and stabilises Ex in the presence of Crb and, remarkably, Ex can be stabilised by either PP2A^Wrd^ or PP2A^Tws^ in the absence of the Crb stimulus. Moreover, the CKI family of protein kinases can phosphorylate Ex in steady-state conditions, and this can be reversed by PP2A. We also demonstrate that Kib forms a complex with PP2A^Wrd^ and Ex and specifically regulates Ex proteostasis. This work provides insight into the crucial role of PP2A as a Hpo signalling regulator, interacting with various upstream components to tightly control the pathway and hence regulate the balance between tissue growth and tissue homeostasis in response to polarity stimuli.

## Materials and Methods

### *Drosophila* cell culture, expression constructs and chemical treatments

*Drosophila* S2 cells (RRID:CVCL_Z992) were cultured in *Drosophila* Schneider’s medium (Gibco) containing 10% (v/v) FBS, 50 μg/mL penicillin, and 50 μg/mL streptomycin. ORFs were PCR amplified from cDNA (*Drosophila* Genomics Resource Center) and cloned into Entry vectors (pDONR-Zeo) using Gateway technology (Thermo Fisher Scientific). FLAG and HA tag expression vectors from the *Drosophila* Gateway Vector Collection and an in-house V5 tag expression vector were used as destination vectors (P. Ribeiro *et al*., 2014). Ex^FL^, Ex^1-468^, Ex^1-468 S453A^, Ex^1-450^, Ex^1-709^ and Ex^710-1427^, Crb^intra^ and Crb^intra ΔFBM^ plasmids were previously described (Genevet *et al*., 2010; Ling *et al*., 2010; Ribeiro *et al*., 2014; Fulford *et al*., 2019). Mts point mutations were generated using the Quikchange Site-Directed Mutagenesis kit (Agilent) according to the manufacturer’s protocol. S2 cells were transfected with Effectene Transfection Reagent (Qiagen) according to the manufacturer’s instructions. Where indicated, proteasome inhibition was achieved by treating cells with 50 μM MG132 (Cambridge Bioscience) and 50 μM calpain inhibitor I (Ac-LLnL-CHO or LLnL) (Sigma) for 4 hr before cell lysis or with 5 μM MG132 overnight.

### RNAi synthesis and treatment

dsRNAs were synthesised using the Megascript T7 kit (Thermo Fisher Scientific) according to the manufacturer’s protocol. DNA templates for dsRNA synthesis were PCR amplified from genomic DNA or plasmids encoding the corresponding genes using primers containing the 5’ T7 RNA polymerase-binding site sequence. The following primers were used:

***lacZ*** fwd: 5’-taatacgactcactataggttgccgggaagctagagtaa-3’;

rev: 5’ taatacgactcactatagggccttcctgtttttgctcac-3’

***wrd*** fwd: 5’-taatacgactcactatagggcgtgagaagctgtcgcaaag-3’;

rev: 5’-taatacgactcactatagggcgctctgatcatacgctgaa-3’

***wdb*** fwd: 5’-taatacgactcactataggggccacgacatcgaacagc-3’;

rev: 5’-taatacgactcactatagggcatttgtggtcgcaggatta-3’

***tws*** fwd: 5’-taatacgactcactatagggtcaacaacttttccagcgtg-3’;

rev: 5’-taatacgactcactatagggtcatttatggtttgcgttttt-3’

***cka*** fwd: 5’-ctaatacgactcactatagggacctggaacgccaagtacac-3’;

rev: 5’-ctaatacgactcactatagggccacagcttaaccgttccat-3’

***mts*** fwd: 5’-ctaatacgactcactatagggtcgactacttgccactgacg;

rev: 5’-ctaatacgactcactataggggcgagcaatccagcttaaagg-3’).

Following cell seeding, S2 cells were incubated with 20 μg dsRNA for 1 hr in serum-free medium and then supplemented with complete medium. Cells were lysed 72 hr after dsRNA treatment and processed as detailed below.

### Immunoprecipitation and immunoblot analysis

For immunoprecipitation of FLAG-tagged proteins, cells were lysed in lysis buffer (50 mM Tris pH 7.5, 150 mM NaCl, 1% Triton X-100, 10% (v/v) glycerol, and 1 mM EDTA) supplemented with phosphatase inhibitor cocktail 2 and 3 (Sigma), and protease inhibitor cocktail (cOmplete Protease Inhibitor, Roche). Cell extracts were spun at 17,000g for 10 min at 4°C. FLAG-tagged proteins were incubated with anti-FLAG M2 Affinity agarose gel (Sigma) for 1-2hrs at 4°C. FLAG immunoprecipitates were washed three to four times with lysis buffer. Elution from beads was performed by incubating with 150 ng/μL 3× FLAG peptide for 15–30 min at 4°C. Detection of purified proteins and associated complexes was performed by immunoblot analysis using chemiluminescence (Immobilon Crescendo/Forte; Thermo Fisher Scientific). Western blots were probed with mouse anti-FLAG (M2; Sigma; RRID:AB_262044), mouse anti-Myc (9E10; Santa Cruz Biotechnology; RRID:AB_262044), rat anti-HA (3F10; Roche Applied Science; RRID:AB_2314622), mouse anti-V5 (Thermo Fisher Scientific; RRID:AB_2556564), or mouse anti-tubulin (E7; DSHB; RRID:AB_528499).

### RNA isolation and RT-PCR analysis

Total RNA was extracted from S2 cells using the QIAshredder and RNeasy kits (Qiagen) according to manufacturer’s instructions. cDNA was synthesised from 1μg of total RNA using the QuantiTect Reverse Transcription kit (Qiagen) as per the manufacturer’s protocol. RT-PCR analysis was performed using 1 μl of cDNA per PCR reaction and the following primers: lacZ (fwd: ttgccgggaagctagagtaa and rev: gccttcctgtttttgctca), tws (fwd: TCAACAACTTTTCCAGCGTG and rev: TCATTTATGGTTTGCGTTTTT). RT-PCR products were run on 2% UltraPure agarose gels (Thermo Fisher Scientific).

### Immunostaining

Larval tissues were processed as previously described (Genevet *et al*., 2010; Ribeiro *et al*., 2014; Fulford *et al*., 2019). Primary and secondary antibodies were incubated overnight at 4°C. Rat anti-Ci (2A1, DSHB; RRID:AB_2109711) was used at 1:100 dilution, mouse anti-β-galactosidase (Z3781, Promega; RRID:AB_430877) was used at 1:500 dilution. Anti-mouse Rhodamine Red-X– conjugated (Jackson ImmunoResearch) was used at 1:250 dilution and anti-rat Alexa 568 (Abcam) was used at 1:200 dilution. After washes, samples were stained with DAPI (1 μg/mL) for 15 min at room temperature before clearing in Vectashield (without DAPI) (H-1200, Vector Labs; RRID:AB_2336790), and mounting with Mowiol 40-88 (Sigma). Fluorescence images were acquired on an inverted Zeiss LSM 880 Airy Scan confocal microscope with a Zeiss Plan-Apochromat 40X/1.3 NA Oil objective (Carl Zeiss).

### *Drosophila* genetics and genotypes

The following transgenic fly stocks were obtained from either the Vienna *Drosophila* Stock Center (VDRC) or Bloomington *Drosophila* Stock Center (BDSC): *UAS-wrd-RNAi* (VDRC 107057KK), *UAS-wdb-RNAi* (VDRC 101406KK), *UAS-tws-RNAi* (VDRC 104167KK), *UAS-Mts* ^BL^ (BDSC 53709). *ubi-Ex*^*1-468*^*::GFP* (Fulford *et al*., 2019), *ubi-Ex*^*1-468*^*S453A::GFP* (Fulford *et al*., 2019), *UAS-Crb* ^*intra*^ (Bulgakova and Knust, 2009), *UAS-lacZ-RNAi* (gift from Nicolas Tapon), HRE-*diap1* ^*GFP4*.*3*^ (gift from Jin Jiang, UT Southwestern) and *fj-lacZ* (Villano and Katz, 1995) have been previously described. The *UAS-Mts* ^*WT*^, *UAS-Mts* ^*H118N*^ and *UAS-Mts* ^*R268A*^ constructs were cloned using Gateway technology into the pUASg-HA(N)-attB vector and transgenic flies were generated by BestGene. Transgenes were inserted at 62E1 (BL-9748) using ΦC31-mediated integration. All crosses were raised at 25°C unless otherwise stated. Genotypes were as follows:

**Figure 1B, 2B, S2B:** *w;; hh-Gal4, UAS-mIFP, ubi-Ex*^*1-468*^ *::GFP*

**Fig 1.**
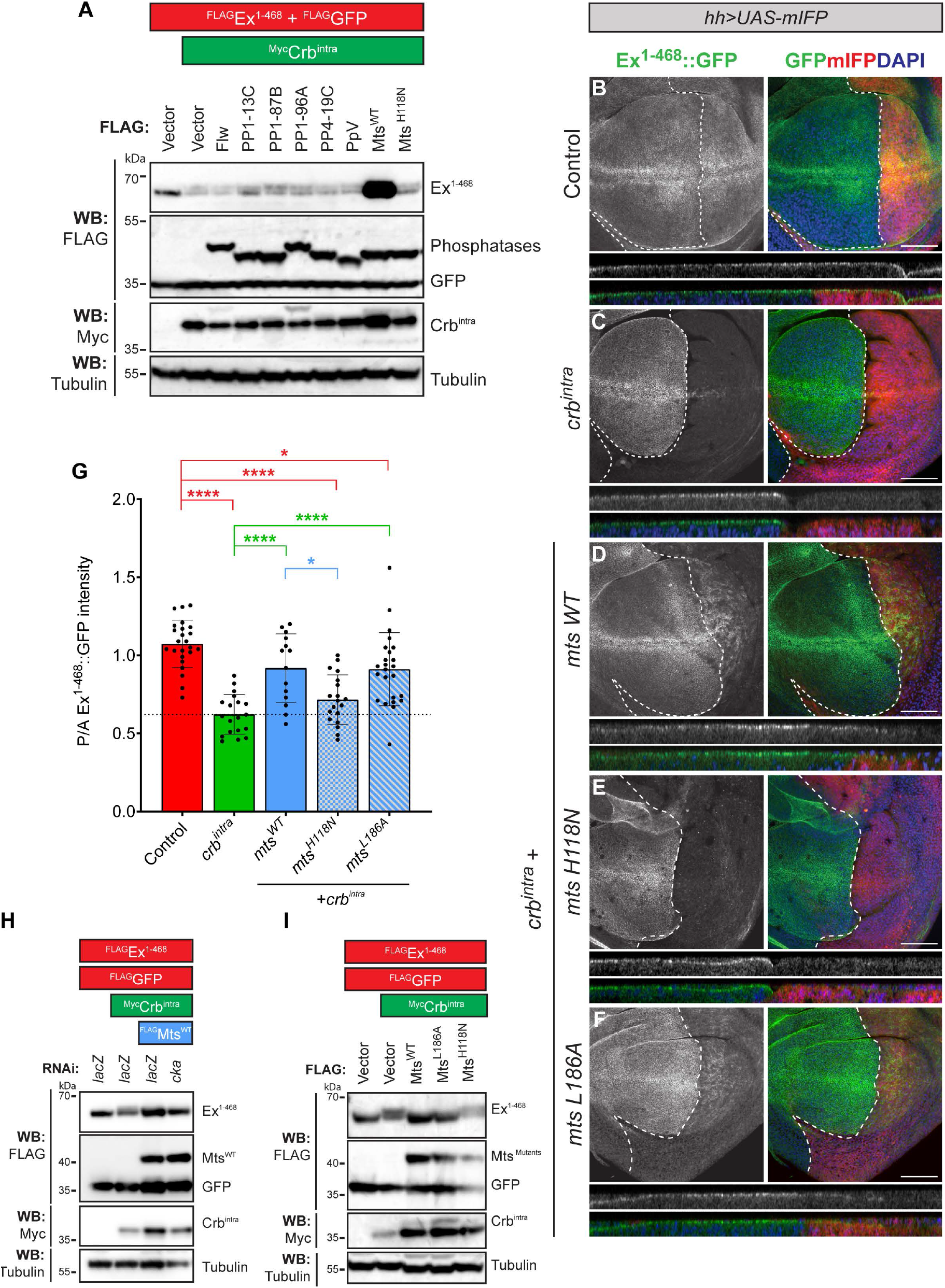
PP2A prevents Crb-induced Ex phosphorylation and degradation. **(A)** PP2A abrogates the effects of Crb^intra^ on Ex protein stability *in vitro*. *Drosophila* S2 cells were transfected with FLAG-tagged Ex^1-468^, GFP, Myc-tagged Crb^intra^, and either of the following FLAG-tagged phosphatase catalytic subunits: Flw, PP1-13C, PP1-87B, PP4-19C, PpV, Mts^WT^ or catalytic Mts mutant, Mts^H118N^. Expression of Crb^intra^ caused a mobility shift and degradation of Ex^1-468^. Note that only Mts^WT^ expression reversed the mobility shift and degradation of Ex^1-468^. **(B-F)** *In vivo* PP2A-mediated Ex stabilisation is independent of Cka function. Confocal micrographs show XY (top) and transverse sections (bottom) of third instar wing imaginal discs expressing *ubi-Ex*^*1-468*^*::GFP* (green) and *UAS-mIFP* (red) alone (control) or in combination with the indicated transgenes expressed in the posterior compartment as listed in individual images, under the control of the *hh-Gal4* driver. Nuclei are stained with DAPI (blue). Dashed white lines depict the AP boundary. Scale bars correspond to 50μm. **(G)** Quantification of *in vivo* Ex^1-468::GFP^ reporter levels. Bar chart depicts the mean ratio of posterior to anterior Ex^1-468^::GFP intensity for the indicated genotypes, with all data points included. Significance was assessed by a one-way ANOVA. Sidak’s multiple comparisons post-hoc test were conducted comparing control, *crb* ^*intra*^ and *mts* ^*WT*^ +*crb* ^*intra*^ conditions against all other genotypes and only significant comparisons are shown. n≥14 for all genotypes. *, p≤0.05; **, p≤0.01; ***, p≤0.001; ****p≤0.0001. Note, Crb^intra^ expression results in degradation of Ex reporter, which can be partially rescued by expression of Mts^WT^, but not Mts^H118N^. Mts^L186A^, a mutant unable to bind Cka, phenocopied Mts^WT^. **(H,I)** *in vitro* PP2A-mediated Ex stabilisation occurs independently of STRIPAK function. (H) S2 cells were treated with either *lacZ dsRNA* or *cka* dsRNA for 24 h, before transfection with indicated constructs. (I) Mts^WT^ and Mts^L186A^ stabilise Ex in the presence of Crb^intra^. S2 cells were transfected with the indicated constructs. 48h after transfection, cells were lysed and lysates were analysed by immunoblot using the indicated antibodies. GFP and Tubulin were used as transfection and loading control, respectively.

**Figure 1C, 3E, S2C, S2G:** *w;; hh-Gal4, UAS-mIFP, ubi-Ex*^*1-468*^ *::GFP/ UAS-crb*^*intra*^

**Figure 1D, 3F:** *w;; hh-Gal4, UAS-mIFP, ubi-Ex*^*1-468*^ *::GFP/ UAS-mts* ^*WT*^, *UAS-crb* ^*intra*^

**Figure 1E:** *w;; hh-Gal4, UAS-mIFP, ubi-Ex*^*1-468*^ *::GFP/ UAS-mts* ^*H118N*^, *UAS-crb* ^*intra*^

**Figure 1F:** *w;; hh-Gal4, UAS-mIFP, ubi-Ex*^*1-468*^*::GFP/ UAS-mts* ^*L186A*^, *UAS-crb* ^*intra*^

**Figure 2C, 4F:** *w;; hh-Gal4, UAS-mIFP, ubi-Ex*^*1-468*^*::GFP/ UAS-mts*^*WT*^

**Fig 2.**
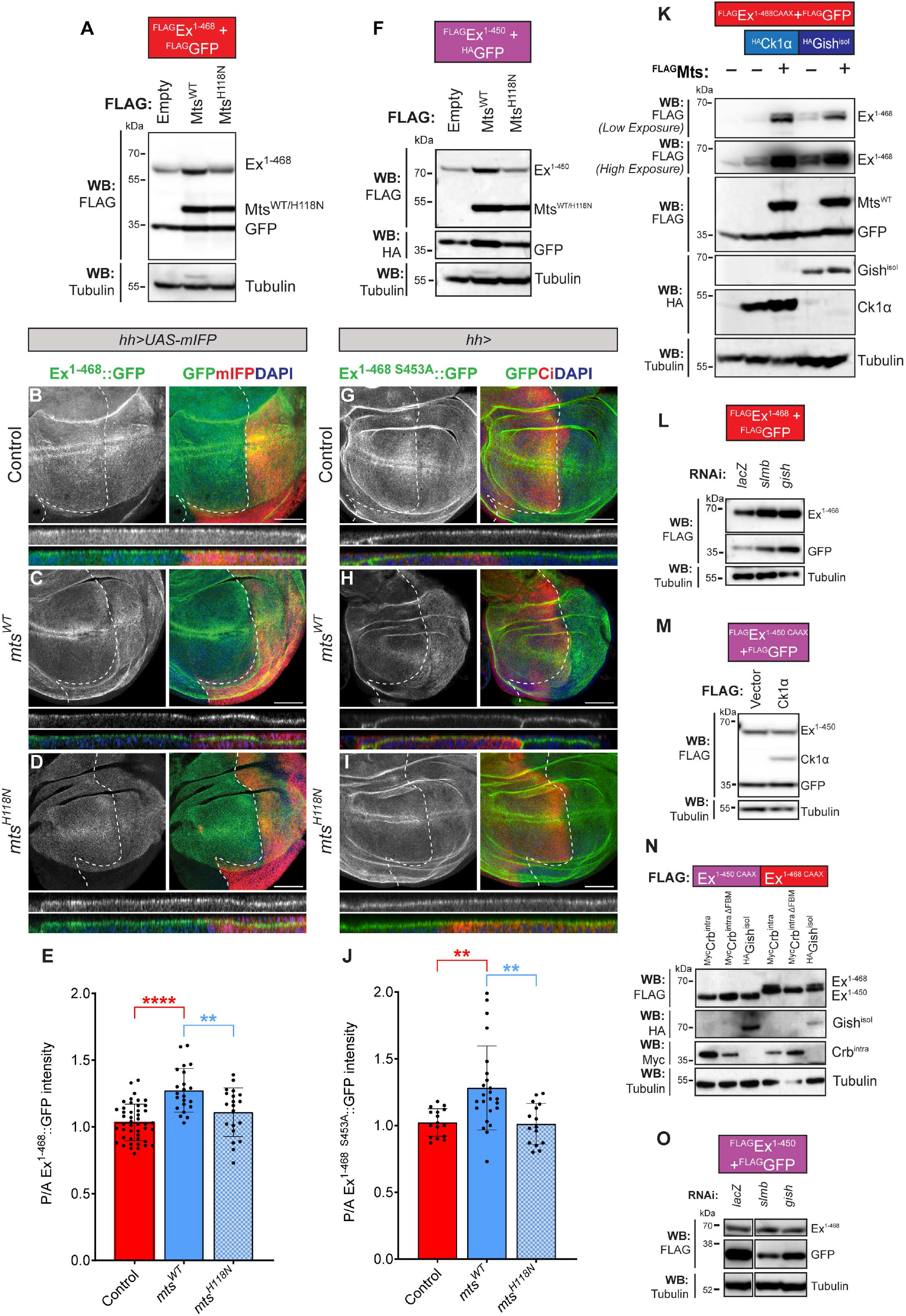
PP2A, CKIs and Slmb regulate steady-state Ex protein levels. **(A)** In the absence of Crb stimulus, Mts^WT^ controls Ex^1-468^ levels in a catalytically-dependent manner. S2 cells were transfected with the indicated constructs and analysed 48h after transfection via immunoblot with the indicated antibodies. **(B-D)** Shown are XY (top) and transverse (bottom) confocal micrographs of third instar wing imaginal discs expressing *ubi-Ex*^*1-468*^*::GFP* (green) and *UAS-mIFP* (red) alone (control, B) or in combination with either *UAS-mts* ^*WT*^ (C) or *UAS-mts* ^*H118N*^ (D) in the posterior compartment, under the control of the *hh-Gal4* driver. Nuclei were stained with DAPI (blue). Dashed white lines depict the AP boundary. Scale bars correspond to 50μm. **(E)** Quantification of Ex^1-468::^GFP reporter levels. Bar chart depicts the mean ratio of posterior to anterior Ex^1-468^::GFP intensity for the indicated genotypes, with all data points included. Significance was assessed by a one-way ANOVA. Tukey’s multiple comparisons post-hoc test were conducted comparing all pairs of genotypes. Only significant comparisons are shown. n≥10 for all genotypes. **, p≤0.01; ****p≤0.0001. Expression of Mts^WT^, but not Mts^H118N^, resulted in an increase in Ex^1-468^ reporter levels in the posterior compartment. **(F)** Mts stabilises Ex independently of the Slmb recognition sequence adjacent to its FERM domain. S2 cells were transfected with FLAG-tagged Ex^1-450^, HA-tagged GFP, and FLAG-tagged Mts^WT^ or Mts^H118N^ and cells were processed 48h after transfection and analysed by immunoblot with the indicated antibodies. Mts stabilised Ex^1-450^ levels in a manner dependent on its catalytic activity. GFP and Tubulin were used as transfection and loading controls, respectively. **(G-I)** Mts controls the protein stability of a Crb-refractory Ex reporter. Confocal micrographs of XY (top) and transverse sections (bottom) of third instar wing imaginal discs expressing *ubi-Ex1-*^*468 S453A*^ *::GFP* (green) alone (control) or in combination indicated transgenes in the posterior compartment, under the control of the *hh-Gal4* driver. Ci staining (red) marks the anterior compartment, where transgenes are not expressed. Nuclei were stained with DAPI (blue). Dashed white lines depict the AP boundary. Scale bars correspond to 50μm. **(J)** Quantification of homeostatic Ex^1-468 S453A::^GFP reporter levels. Bar chart depicts the mean ratio of posterior to anterior Ex^1-468 S453A^::GFP intensity for the indicated genotypes, with all data points included. Significance was assessed by a Kruskal-Wallis test. Dunn’s multiple comparisons post-hoc test were conducted comparing all pairs of genotypes. Only significant comparisons are shown. n≥15 for all genotypes. **, p<0.01. Expression of Mts^WT^, but not Mts^H118N^, resulted in an increase in Ex^1-468 S453A^ reporter levels in the posterior compartment. **(K)** CKI kinase-mediated Ex phosphorylation is abrogated by PP2A. S2 cells were transfected with the indicated plasmids and analysed 48h after transfection by immunoblot with the indicated antibodies. Ck1α or Gish^isoI^ phosphorylated Ex^1-468 CAAX^, and Mts^WT^ co-expression of either kinase with reversed this phosphorylation and stabilised Ex^1-468 CAAX^. **(L-O)** CKI kinases and Slmb regulate steady-state Ex levels via the Ex^452-457^ Slmb recognition site. (L, O) S2 cells were treated with dsRNA for 24h before transfecting with the indicated constructs. In all cases, cells were lysed and processed for immunoblot analysis with the indicated antibodies 48h after transfection. (L) Knocking down either *slmb, gish* or *ck1α* resulted in an increase in homeostatic Ex^1-468^ levels in the absence of ectopic Crb. (M) Ck1α is unable to phosphorylate Ex^1-450^. (N) Expression of Crb^intra ΔFBM^ captures homeostatic Ex levels, since Crb^intra^ requires the FBM to promote Ex^1-468^ phosphorylation. Expression of Crb^intra ΔFBM^ does not affect phosphorylation status of either Ex^1-468^ or Ex^1-450^. Expression of Gish^isoI^ promotes phosphorylation of Ex^1-468^ but not Ex^1-450^. (O) RNAi-mediated depletion of either *slmb* or *gish* did not affect Ex^1-450^ levels. GFP and Tubulin were used as transfection or loading controls, respectively.

**Figure 2D:** *w;; hh-Gal4, UAS-mIFP, ubi-Ex*^*1-468*^*::GFP/ UAS-mts* ^*H118N*^

**Figure 2G:** *w;; hh-Gal4, ubi-Ex1-*^*468 S453A*^ *::GFP*

**Figure 2H:** *w;; hh-Gal4, UAS-mIFP, ubi-Ex*^*1-468 S453A*^ *::GFP / UAS-mts* ^*WT*^

**Figure 2I:** *w;; hh-Gal4, UAS-mIFP, ubi-Ex*^*1-468 S453A*^*::GFP / UAS-mts* ^*H118N*^

**Figure 3D, 4A, S2F:** *w;; hh-Gal4, UAS-mIFP, ubi-Ex*^*1-468*^ *::GFP / UAS-lacZ-RNAi*

**Fig 3.**
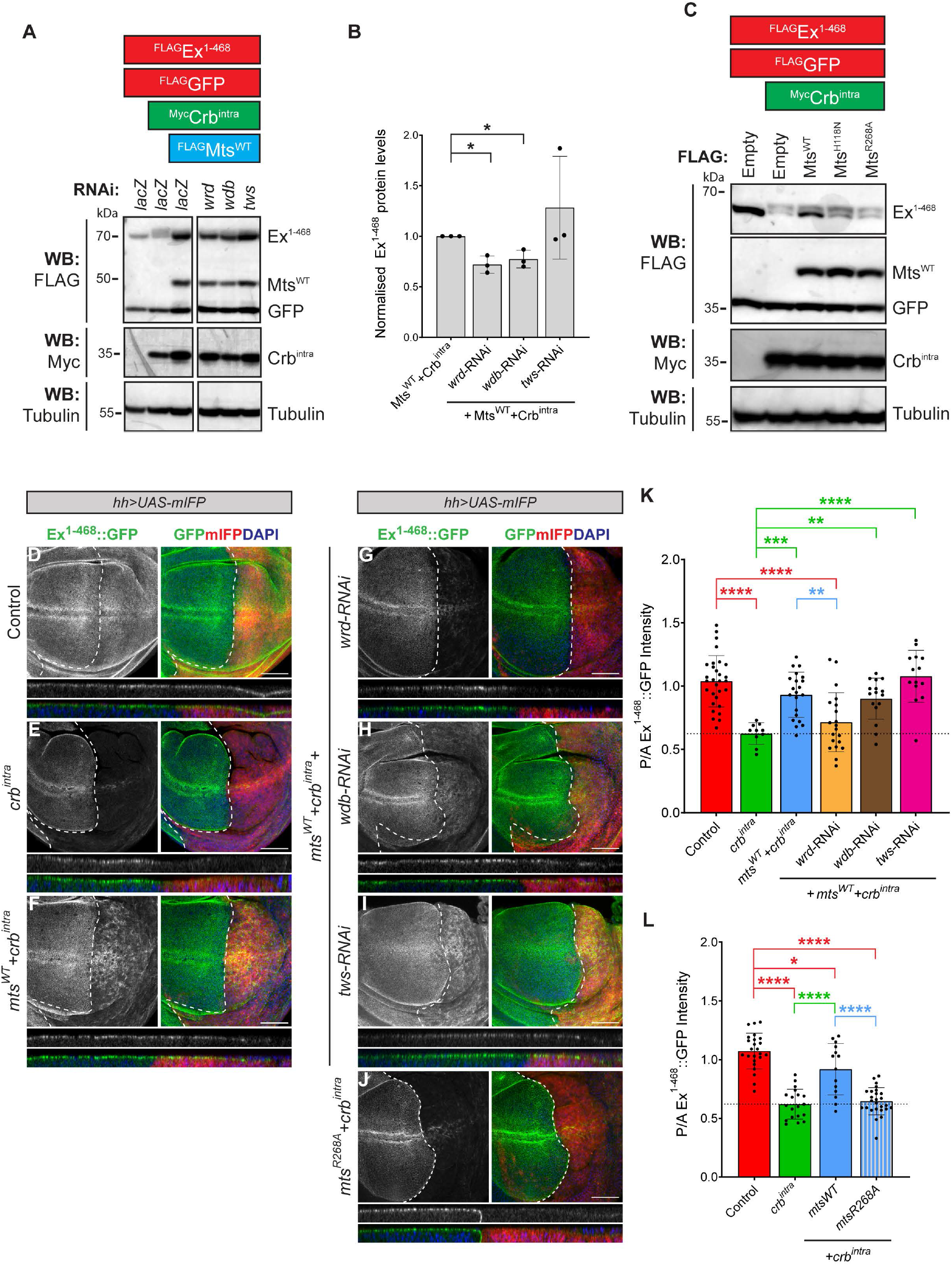
Wrd is required for PP2A-dependent stabilisation of Ex in the presence of Crb. **(A)** S2 cells were treated with either *wrd, wdb* or *tws* dsRNA for 24 h before transfection with the indicated constructs. Cells were lysed and analysed by immunoblot using the indicated antibodies 48h after transfection. GFP and Tubulin were used as transfection or loading controls, respectively. **(B)** Quantification of *in vitro* Ex^1-468^ protein levels. Bar chart depicts normalised Ex^1-468^ levels (to the corresponding Mts^WT^+Crb^intra^ sample) from 3 independent experiments. Significance was assessed by an unpaired t-test between Mts^WT^+Crb^intra^ and other conditions and only significant comparisons are shown. *, p<0.05. **(C)** PP2A^Wrd^ regulates Ex stability. S2 cells were transfected with the indicated constructs 48h before lysis and Western blot analysis with the indicated antibodies. Note that Mts^R268A^, a mutant that affects binding to Wrd, was unable to stabilise Ex^1-468^ in the presence of Crb^intra^. GFP and Tubulin were used as transfection and loading controls, respectively. **(D-J)** *In vivo* assessment of the role of PP2A regulatory subunits in the regulation of the Ex reporter. Confocal micrographs depict XY (top) and transverse sections (bottom) of third instar wing imaginal discs expressing *ubi-Ex*^*1-468*^ *::GFP* (green) and *UAS-mIFP* (red) alone (control, D) or in combination with indicated transgenes in the posterior compartment, under the control of the *hh-Gal4* driver. Nuclei were stained with DAPI (blue). Dashed white lines depict the AP boundary. Scale bars correspond to 50μm. **(K,L)** Quantification of effect of PP2A regulato
ry subunits on Ex *in vivo* reporter levels. Bar chart depicts the mean ratio of posterior to anterior Ex^1-468^::GFP intensity for the indicated genotypes, with all data points represented. Significance was assessed by a one-way ANOVA. Sidak’s multiple comparisons post-hoc test were conducted comparing control, *crb* ^*intra*^ and *mts* ^*WT*^ +*crb* ^*intra*^ conditions against all other genotypes and only significant comparisons are shown.n≥10 for all genotypes. *, p<0.05; **, p<0.01; ***, p<0.001; ****p<0.0001. wrd depletion abrogates the ability of Mts^WT^ to stabilise Ex in the presence of Crb^intra^, while mts^R268A^ expression does not stabilise Ex in the presence of Crb^intra^.

**Figure 3G:** *w; UAS-wrd-RNAi (VDRC 107057KK); hh-Gal4, UAS-mIFP, ubi-Ex*^*1-468*^ *::GFP / UAS-mts*^*WT*^, *UAS-Crb*^*intra*^

**Figure 3H:** *w; UAS-wdb-RNAi (VDRC 101406KK); hh-Gal4, UAS-mIFP, ubi-Ex*^*1-468*^ *::GFP / UAS-mts* ^*WT*^, *UAS-Crb* ^*intra*^

**Figure 3I:** *w; UAS-tws-RNAi (VDRC 104167KK); hh-Gal4, UAS-mIFP, ubi-Ex^1-468^::GFP / UAS-mts ^WT^, UAS-Crb ^intra^*D

**Figure 3J:** *w;; hh-Gal4, UAS-mIFP, ubi-Ex*^*1-468*^*::GFP / UAS-mts* ^*R268A*^, *UAS-Crb* ^*intra*^

**Figure 4B:** *w;; hh-Gal4, UAS-mIFP, ubi-Ex*^*1-468*^*::GFP / UAS-mts* ^*BL*^ *(BL 53709)*

**Fig 4.**
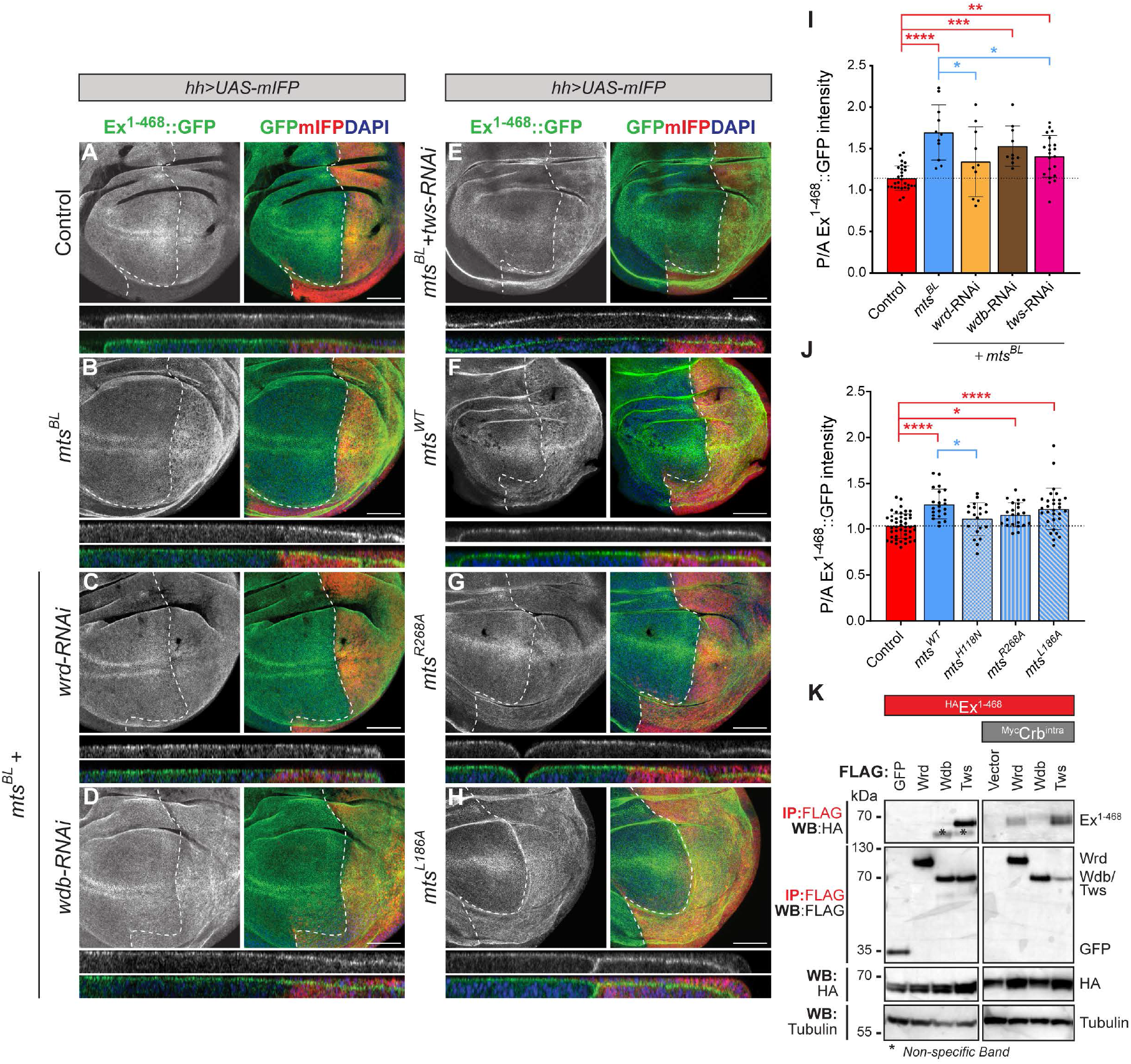
PP2A ^Wrd^ and PP2A ^Tws^ are involved in regulation of Ex proteostasis. **(A-H)** Confocal micrographs show XY (top) and transverse sections (bottom) of third instar wing imaginal discs expressing *ubi-Ex*^*1-468*^ *::GFP* (green) and *UAS-mIFP* (red) alone (control, A) or in combination with indicated transgenes in the posterior compartment, under the control of the *hh-Gal4* driver. Nuclei were stained with DAPI (blue). Dashed white lines depict the AP boundary. Scale bars correspond to 50μm. **(I,J)** Quantification of Ex reporter levels. Bar chart depicts the mean ratio of posterior to anterior Ex^1-468^::GFP intensity for the indicated genotypes, with all data points depicted. Significance was assessed by a one-way ANOVA. Sidak’s multiple comparisons post-hoc test were conducted comparing control and *mts* ^*BL*^ conditions against all other genotypes and only significant comparisons are shown. n≥10 for all genotypes. *, p<0.05; **, p<0.01; ***, p<0.001; ****p<0.0001. Note that knocking down wrd abrogates the ability of Mts^WT^ to increase Ex levels in the absence of Crb, while overexpression of mts^R268A^ does not increase Ex levels. **(K)** Ex interaction with PP2A regulatory subunits. S2 cells were transfected with the indicated constructs. For conditions where Myc-tagged Crb^intra^ was transfected, cells were treated with MG132 to avoid Ex degradation. Following lysis, FLAG-tagged proteins were purified using FLAG-agarose beads. Lysates and eluates were processed for Western blotting analysis with the indicated antibodies. In the absence of Crb^intra^, Ex co-immunoprecipitated with Tws; in the presence of Crb^intra^, Ex co-immunoprecipitated with Wrd and Tws. Tubulin was used as protein loading control.

**Figure 4C:** *w; UAS-wrd-RNAi (VDRC 107057KK); hh-Gal4,UAS-mIFP, ubi-Ex*^*1-468*^*::GFP / UAS-mts* ^*BL*^ *(BL 53709)*

**Figure 4D:** *w; UAS-wdb-RNAi (VDRC 101406KK); hh-Gal4,UAS-mIFP, ubi-Ex*^*1-468*^ *::GFP / UAS-mts* ^*BL*^ *(BL 53709)*

**Figure 4E:** *w; UAS-tws-RNAi (VDRC 104167KK); hh-Gal4, UAS-mIFP, ubi-Ex*^*1-468*^ *::GFP / UAS-mts* ^*BL*^ *(BL 53709)*

**Figure 4G:** *w;; hh-Gal4, UAS-mIFP, ubi-Ex*^*1-468*^*::GFP / UAS-mts* ^*R268A*^

**Figure 4H:** *w;; hh-Gal4, UAS-mIFP, ubi-Ex*^*1-468*^ *::GFP / UAS-mts* ^*L186A*^

**Figure 6A:** *w; en-Gal4, UAS-RFP; HRE-diap1* ^*GFP4*.*3*^

**Figure 6B:** *w; en-Gal4, UAS-RFP; HRE-diap1* ^*GFP4*.*3*^ */ UAS-crb* ^*intra*^

**Figure 6C:** *w; en-Gal4, UAS-RFP; HRE-diap1* ^*GFP4*.*3*^*/ UAS-mts* ^*WT*^, *UAS-crb* ^*intra*^

**Figure 6D:** *w; en-Gal4, UAS-RFP; HRE-diap1* ^*GFP4*.*3*^ */ UAS-mts* ^*H118N*^, *UAS-crb* ^*intra*^

**Figure 6E:** *w; en-Gal4, UAS-RFP; HRE-diap1*^*GFP4*.*3*^*/ UAS-mts*^*L186A*^, *UAS-crb*^*intra*^

**Figure 6F:** *w; en-Gal4, UAS-RFP; HRE-diap1* ^*GFP4*.*3*^ */ UAS-mts* ^*R268A*^, *UAS-crb* ^*intra*^

**Figure S2D, S2H:** *w;; hh-Gal4, UAS-mIFP, ubi-Ex*^*1-468*^*::GFP/ mts* ^*BL*^ *(BL 53709), UAS-crb* ^*intra*^

**Figure S2I:** *w; UAS-cka-RNAi (VDRC 104167KK); hh-Gal4, UAS-mIFP, ubi-Ex*^*1-468*^ *::GFP/ UAS-mts* ^*BL*^ *(BL 53709), UAS-crb* ^*intra*^

**Figure S5A:** *w;; hh-Gal4, UAS-CD8::GFP, fj-lacZ*

**Figure S5B:** *w;; hh-Gal4, UAS-CD8::GFP, fj-lacZ / UAS-crb* ^*intra*^

**Figure S5C:** *w;; hh-Gal4, UAS-CD8::GFP, fj-lacZ / UAS-mts* ^*WT*^, *UAS-crb* ^*intra*^

**Figure S5D:** *w;; hh-Gal4, UAS-CD8::GFP, fj-lacZ / UAS-mts* ^*H118N*^, *UAS-crb* ^*intra*^

**Figure S5E:** *w;; hh-Gal4, UAS-CD8::GFP, fj-lacZ / UAS-mts* ^*L186A*^, *UAS-crb* ^*intra*^

### Immunofluorescence quantification and statistical analyses

To assess changes in Ex^1-468^::GFP or Ex^1-468 S453A^::GFP reporter levels, the ratio of apical GFP intensity in the posterior versus anterior compartment in two transverse sections per wing disc was measured using Fiji. For the Hpo activity reporter experiments using *diap1* ^*GFP4*.*3*^, the sum of all Z-stack confocal slices were projected using Fiji and the ratio of total intensity of the reporter in posterior versus anterior compartments was measured using Fiji. For all experiments, mean, geometric mean or rank of the posterior:anterior intensities were compared across different genotypes. The normality or lognormality of the data was determined using the Shapiro-Wilk, D’Augostino and Pearson, Anderson-Darling and Kolmogorov-Smirnov normality tests. Bartlett’s homogeneity of variance test was used to check for similarity of variances. If data was normally distributed and variances were similar, one-way ANOVA followed by Tukey’s or Sidak’s post-hoc test were used and if variances were dissimilar, Welch ANOVA followed by Dunnett’s T3 post-hoc test was used. If data was log-normally distributed, the above analysis pipeline was used on log-transformed data. If data was not normally distributed, the ranks were compared across different conditions using Kruskal-Wallis test followed by Dunn’s post-hoc test. When the data were log-transformed, the analyses conducted compared the geometric means of the data rather than the arithmetic means. For densitometry analyses conducted in Fig. 3B, Ex protein levels were normalised to levels in Mts^WT^+Crb^intra^, and multiple paired t-tests were conducted. For comparing binding efficiencies of Mts mutants to corresponding regulatory subunits (Fig. S3H, S5D-E) paired t-tests were conducted. Differences between experimental groups were deemed significant if the p-value was < 0.05. Given the gradient expression pattern of *fj-lacZ*, we utilised the Fire lookup table (LUT) in Fiji to clearly visualise the changes in intensity levels of the reporter.

## Results

### PP2A antagonises Crb-mediated Ex phosphorylation and degradation

In addition to recruiting Ex to the apical membrane to activate the Hippo pathway, Crb also regulates Ex turnover by promoting Ex phosphorylation by Ck1α and Gish, which subsequently results in Ex ubiquitylation and degradation by a Slmb-containing ubiquitin ligase complex (Ribeiro *et al*., 2014; Fulford *et al*., 2019). As the dynamic regulation of Ex is a crucial determinant of Hpo pathway activity, we hypothesised that the Ex degradation promoted by Crb would be balanced by a counteracting mechanism that can reverse Crb-mediated Ex phosphorylation, thereby stabilising Ex. To elucidate this mechanism, we tested a panel of serine/threonine protein phosphatases for their ability to dephosphorylate and stabilise Ex in the presence of Crb. Expression of either full-length (Crb^FL^) or the intracellular domain of Crb (Crb^intra^) in *Drosophila* S2 cells promotes delayed electrophoretic mobility of Ex (indicative of Ex phosphorylation) and reduced Ex protein levels (Ribeiro *et al*., 2014; Fig. 1A). Moreover, Ex^1-468^, which comprises the N-terminal FERM domain and the consensus Slmb recognition site ^452^TSGIVS^457^, is the minimal region required for Crb-mediated Ex phosphorylation and degradation (Ribeiro *et al*., 2014). We co-expressed Ex^1-468^ and the catalytic subunits of phosphatases PP1 (Flapwing (Flw), PP1-13C and PP1-87B), PP4 (PP4-19C), PP6 (PpV) and PP2A (Microtubule star (Mts)), in the presence of Crb^intra^ in S2 cells to test if any of the phosphatases could prevent Crb-mediated Ex phosphorylation and degradation (Fig.1A). Strikingly, we observed that, contrary to the catalytic subunits of the other phosphatases, Mts prevented the Ex mobility shift induced by Crb^intra^ and stabilised Ex. Since Mts shares 98.4% protein sequence similarity with its human counterpart (PPP2CA; Fig. S1), we generated a Mts catalytic mutant (Mts^H118N^) based on existing PPP2CA structural data (Evans *et al*., 1999), to eliminate potential pleiotropic effects associated with Mts overexpression. We validated the mutant’s lack of catalytic activity by demonstrating that Mts^H118N^, unlike wild-type Mts (hereafter referred to as Mts^WT^), was unable to stabilise Armadillo (Arm, the *Drosophila* β-catenin orthologue), a previously characterised PP2A substrate (Bajpai *et al*., 2004), in S2 cells (Fig. S2A). We then tested the effect of Mts^H118N^ on Crb-mediated Ex phosphorylation and degradation and observed that Mts^H118N^ was unable to stabilise Ex in the presence of Crb^intra^ (Fig. 1A). Furthermore, Mts restricted Crb-mediated Ex degradation in a dose-dependent manner, which suggests a dynamic competition between Crb- and PP2A-dependent regulation of Ex (Fig. S2B). Therefore, our results demonstrate that PP2A counteracts Ex phosphorylation and degradation promoted by Crb.

Next, we tested whether PP2A antagonises Crb-mediated Ex degradation *in vivo*. Since ex is a Yki target gene, we used an *in vivo* Ex protein stability reporter, *ubi-Ex*^*1-468*^ *::GFP* (Fulford *et al*., 2019) to specifically study post-translational regulation of Ex *in vivo*. This reporter expresses GFP-tagged Ex^1-468^ under the control of the *ubiquitin 63E* promoter rather than the endogenous ex promoter, thus decoupling the effect of transcriptional feedback from the Hpo pathway on Ex protein levels and mitigating any confounding effects associated with Ex-dependent regulation of the Hpo pathway (Fulford *et al*., 2019). Transgenes were expressed in the posterior compartment of the larval wing imaginal disc using *hedhehog-Gal4* (*hh-Gal4*). The ratio of mean apical Ex^1-468^::GFP intensity between the posterior and the anterior compartments was used as a measure of normalised Ex protein levels. Expression of Crb^intra^ resulted in a prominent reduction of apical Ex^1-468^::GFP levels in the posterior compartment, recapitulating the Crb-mediated degradation of Ex observed *in vitro* (Fig. 1B,C,G). To study if PP2A regulates Crb-mediated Ex degradation *in vivo*, we generated transgenic flies carrying either an *UAS-mts*^*WT*^ or a *UAS-mts*^*H118N*^ construct. We observed that expression of Mts^WT^ suppressed Crb^intra^-mediated reduction in apical Ex protein levels (Fig. 1B,D,G,I). We validated these results using an alternative *UAS-mts* stock (obtained from the Bloomington stock collection and hereafter referred to as *UAS-mts*^*BL*^; Fig. S2C-F). In contrast, Mts^H118N^ was unable to stabilise Ex in the presence of Crb^intra^ (Fig. 1B,E,G). Together, our data suggest that PP2A dephosphorylates Ex and prevents its degradation in the presence of Crb.

### PP2A stabilises Ex independently of the STRIPAK complex ^WT^

We and others have delineated a crucial role for the STRIPAK complex in the regulation of Hpo signalling (Ribeiro *et al*., 2010; Bae *et al*., 2017; Zheng *et al*., 2017). The STRIPAK complex is a large multicomponent protein complex that contains PP2A and, within it, PP2A substrate specificity is conferred by the Striatin regulatory subunit, Cka, which leads to Hpo dephosphorylation and Hpo pathway inactivation (Ribeiro *et al*., 2010; Jeong *et al*., 2021). Hence, we tested whether PP2A-mediated regulation of Ex involved the STRIPAK complex. RNAi-mediated depletion of *cka* in S2 cells did not hinder the ability of Mts to stabilise Ex^1-468^ in presence of Crb^intra^, suggesting that the effect of PP2A is independent of the STRIPAK complex (Fig.1H). To test whether Cka is part of the PP2A complex that antagonises Crb-mediated Ex degradation *in vivo*, we knocked down cka while simultaneously overexpressing Mts (*UAS-mts* ^*BL*^) and Crb^intra^ in the posterior compartment of the larval wing disc using *hh-Gal4* (Fig. S3A-D). Since Cka is required for many developmental processes (Chen *et al*., 2002; La Marca *et al*., 2019), crosses were raised at 18°C to avoid developmental defects in the posterior compartment of the larval wing disc, which were consistently observed at 25°C (data not shown). However, in these conditions, we were unable to observe significant Crb-mediated degradation of Ex^1-468^::GFP (Fig.S3E). Hence, using this approach we were unable to determine whether Cka is required for PP2A-mediated regulation of Ex protein levels *in vivo*.

To overcome this limitation and dissect whether the STRIPAK complex is involved in Ex stabilisation *in vivo*, we identified and mutated a conserved residue in Mts (Mts^L186A^; Fig. S1) that was shown to affect binding of PPP2CA with STRN3, the mammalian orthologue of Cka (Jeong *et al*., 2021). We validated that Mts^L186A^ displays impaired binding to Cka compared to Mts^WT^ by performing co-immunoprecipitation (co-IP) experiments in S2 cells (Fig. S3F,G). Additionally, using the Arm stabilisation assay described above, we determined that the L186A mutation does not hinder the catalytic activity of Mts (Fig. S3H). In contrast to Mts^H118N^, expression of Mts^L186A^ in S2 cells did not affect the ability of Mts to dephosphorylate and stabilise Ex in the presence of Crb^intra^ (Fig. 1H). Accordingly, *hh-Gal4*-mediated co-expression of Mts^L186A^ with Crb^intra^ in the posterior compartment resulted in increased apical Ex protein levels when compared with those discs expressing only Crb^intra^ (Fig. 1B,C,F,I). Therefore, like Mts^WT^, Mts^L186A^ can partially rescue Ex in the presence of Crb^intra^. These results suggest that the STRIPAK complex is not essential for limiting Crb-mediated Ex regulation.

### PP2A regulates steady-state Ex protein levels via a mechanism distinct from Crb-dependent control

While studying how PP2A regulates Ex in the context of Crb expression, we also observed that PP2A regulated Ex stability in the absence of ectopic Crb expression. Overexpression of Mts^WT^ in the absence of the Crb^intra^ stimulus resulted in increased Ex protein levels in S2 cells (Fig. 2A), while the expression of the Mts catalytic mutant, Mts^H118N^, did not have any effect on steady-state Ex protein levels (Fig. 2A). Moreover, knocking down mts resulted in a decrease in Ex protein levels in S2 cells (Fig. S4A). In sharp contrast, overexpression of the catalytic subunits of PP1s, PP4 and PP6 did not affect Ex protein levels in the absence of Crb^intra^ expression (Fig. S4B). Given the extremely low expression in S2 cells, these results suggest that PP2A can also stabilise Ex in a manner independent of Crb function. We then extended our analysis to *in vivo* conditions, where expression of Mts^WT^ in the posterior compartment of the wing disc using hh-Gal4 driver resulted in increased levels of the *ubi-Ex*^*1-468*^ *::GFP* reporter (Fig. 2B,C,E). In contrast, expression of Mts^H118N^ had no effect on steady-state Ex levels (Fig. 2B,D,E). These results suggest that PP2A is essential for maintenance of Ex protein levels, a process we refer to as regulation of Ex proteostasis. Since the catalytic activity of PP2A is essential to stabilise Ex, this suggests that Ex is phosphorylated even in the absence of Crb expression.

Given that previous studies have demonstrated that the N-terminal and C-terminal regions of Ex are differentially controlled (Ribeiro *et al*., 2014; Ma *et al*., 2018; Fulford *et al*., 2019; Song and Ma, 2023), we tested whether PP2A can regulate the stability of the Ex truncation mutants, Ex^469-1030^ and Ex^1031-1427^ (Ribeiro *et al*., 2014). Expression of Mts^WT^ did not modulate the protein levels of either Ex^469-1030^ or Ex^1031-1427^ (Fig. S4C). Furthermore, as expected, we verified that PP2A was able to stabilise full-length Ex (Fig. S4D). Therefore, in conditions where ectopic Crb is absent, PP2A stabilises Ex exclusively by interacting with the N-terminal region of Ex. We previously described a crucial role for the Slmb recognition site Ex^452-457^ in Crb-mediated Ex phosphorylation and degradation (Ribeiro *et al*., 2014). Hence, we investigated whether this Slmb degron was also involved in regulation of Ex proteostasis in S2 cells. For this, we tested whether Mts^WT^ can stabilise Ex^1-450^, a truncation lacking the Slmb degron and, therefore, refractory to Crb regulation (Ribeiro *et al*., 2014). Interestingly, we observed that Mts^WT^ was able to stabilise Ex^1-450^ and, more importantly, this PP2A-mediated stabilisation of Ex^1-450^ required its catalytic activity (Fig. 2F). To investigate whether this regulatory mechanism requires the 452-457 Slmb degron *in vivo*, we used the *ubi-Ex*^*1-468 S453A*^ *::GFP* reporter, which is mutated within the Slmb recognition site (Fulford *et al*., 2019). Expression of Mts^WT^, but not Mts^H118N^, in the posterior compartment of the wing disc resulted in an increase in the levels of the *ubi-Ex* ^*1-468 S453A::GFP*^reporter (Fig. 2G-J). This corroborates our *in vitro* observations, indicating that PP2A-mediated regulation of Ex proteostasis is independent of the S453 phosphodegron. These results suggest that Ex is likely phosphorylated at additional sites distinct from the S453 site that is implicated in Crb-mediated Ex regulation. However, whether the S453 site is also phosphorylated in these conditions and whether PP2A can regulate Ex via this site was unclear. Therefore, in addition to the previously characterised Crb-dependent regulation of Ex, a more intricate phosphorylation-dependent mechanism is involved in the maintenance of Ex protein levels.

### CKIs and Slmb regulate Ex proteostasis via the 452-457 Slmb consensus sequence

In the presence of Crb, the CKI family kinases Gish and Ck1α promote phosphorylation of Ex, which results in its Slmb-dependent protein degradation (Ribeiro *et al*., 2014; Fulford *et al*., 2019). Since we observed that, in the absence of the Crb stimulus, PP2A-mediated regulation of Ex occurs at sites other than the Ex^452-457^ Slmb recognition site, we sought to determine whether Gish and Ck1α were involved in this mechanism. The effects of Gish and CK1α on Ex^1-468^ phosphorylation require its membrane localisation (Fulford *et al*., 2019) and, hence, a C-terminal CAAX tag (Sotillos *et al*., 2004) was added to target Ex to the membrane (Ex^1-468 CAAX^) (Fulford *et al*., 2019), allowing us to assess the effect of the kinases in the absence of Crb. Expression of either Gish or CK1α was able to promote Ex^1-468 CAAX^ phosphorylation and this was reversed by expression of Mts^WT^ (Fig. 2K). Furthermore, RNAi-mediated depletion of gish in S2 cells resulted in an increase in Ex^1-468^ levels in the absence of Crb^intra^ (Fig. 2L), suggesting that CKIs may also be involved in regulation of Ex protein levels independently of the action of Crb. Since Gish and Ck1α act via the 452-457 phosphodegron site in the presence of the Crb stimulus (Fulford *et al*., 2019), we next assessed how CKIs regulate Ex proteostasis. We overexpressed either Gish or Ck1α and tested whether they phosphorylate Ex^1-450 CAAX^, a truncation lacking the 452-457 phosphodegron region. Interestingly, we observed that Ex^1-450 CAAX^ was not phosphorylated by Gish or Ck1α (Fig. 2M,N). Additionally, knocking down gish did not alter Ex^1-450^ protein levels, in contrast to the previously observed increased Ex^1-468^ protein levels (Fig. 2O).

Therefore, unlike PP2A, in the absence of Crb function, the regulation of Ex protein levels by Gish and Ck1α appears to be limited to the Ex^452-457^ site.

Since phosphorylation of Ex mediated by Gish and Ck1α promotes Slmb-dependent ubiquitylation and degradation, we wanted to determine whether Crb-independent regulation of Ex protein stability is also controlled by Slmb. RNAi-mediated knockdown of Slmb resulted in an increase in Ex^1-468^ protein levels (Fig. 2L), suggesting that Slmb is indeed involved in regulation of Ex proteostasis. However, we observed that knocking down Slmb did not affect Ex^1-450^ protein levels (Fig. 2O). These results suggest that Slmb regulates the N-terminus of Ex specifically via the 452-457 phosphodegron site. Taken together, these results show that the machinery that facilitates Crb-mediated Ex phosphorylation and degradation is also partly involved in the Crb-independent regulation of Ex protein stability. ^Wrd^

### PP2A^Wrd^ antagonises Crb-mediated Ex degradation while both PP2A^Wrd^ and PP2A^Tws^ regulate Ex at steady-state

As the substrate specificity of the PP2A holoenzyme is defined by its regulatory subunit, we aimed to identify the regulatory subunit(s) involved in the PP2A-mediated mechanism that counteracts Crb-mediated Ex degradation. We generated and validated dsRNAs for the B regulatory subunits, Wrd and Wdb, and the B’ regulatory subunit, Tws (Fig. S5A,B). Given that the regulatory subunits CG32568 (B) and CG4733 (B’’) are not expressed in S2 cells where PP2A can stabilise Ex in presence of Crb, this suggests that these subunits are not involved in Ex regulation and, therefore, they were not tested further. To identify the PP2A regulatory subunit(s) involved in Ex regulation, we depleted *wrd, wdb* or *tws* in conjunction with Mts^WT^ and Crb^intra^ expression in S2 cells (Fig. 3A,B). In these conditions, depletion of a PP2A regulatory subunit involved in this process would be expected to suppress the effect of Mts on Ex protein levels. We observed that when either *wrd* or *wdb* were knocked down, Mts was unable to dephosphorylate and stabilise Ex to the same extent as in control conditions. This suggests that PP2A holoenzymes comprised of either Wrd or Wdb as regulatory subunits antagonise Crb-mediated Ex phosphorylation and degradation. To test whether these regulatory subunits are necessary *in vivo*, we knocked down *wrd, wdb* or *tws* while simultaneously co-expressing Mts^WT^ and Crb^intra^ in the posterior compartment of the wing disc using *hh-Gal4* (Fig. 3D-I,K). RNAi-mediated depletion of *wrd*, in the genetic background of *UAS-mts*^*WT*^ *+UAS-crb* ^*intra*^, inhibited the ability of Mts^WT^ to stabilise the levels of the Ex^1-468^::GFP reporter in the presence of Crb^intra^ (Fig. 3G,K). In sharp contrast, knocking down either *wdb* or *tws* had no effect (Fig. 3H-K), suggesting that PP2A^Wrd^ might play a more prominent role in regulating Ex levels *in vivo*.

Since Wrd and Wdb can be functionally redundant, as they belong to the same PP2A regulatory subunit family (Chen *et al*., 2007; Pinto and Orr-Weaver, 2017; Jang *et al*., 2021), we performed an orthogonal approach to validate the involvement of Wrd in mediating Ex stability *in vivo*. We mutated a conserved residue in Mts (Mts^R268A^; Fig. S1), which was predicted to affect binding of mammalian PPP2CA with its corresponding B regulatory subunits, according to structural biology studies (Cho and Xu, 2006; Xu *et al*., 2006). Using co-IP experiments, we tested whether the mutations indeed affect binding to Wrd and Wdb as predicted. Surprisingly, we found that Mts^R268A^ specifically affected binding to Wrd, but not to Wdb (Fig. S5C-E). Furthermore, using the Arm stabilisation assay, we showed that Mts^R268A^ retains its catalytic activity (Fig S2A). When Mts^R268A^ was co-expressed with Crb^intra^ in S2 cells, we found that Mts^R268A^ was unable to dephosphorylate and stabilise Ex (Fig. 3C). Similar results were observed *in vivo*, where co-expression of Mts^R268A^ and Crb^intra^ in the posterior compartment of the wing disc did not alter the levels of the Ex reporter when compared with expression of Crb^intra^ alone (Fig. 3D-F,J,L). Therefore, we conclude that Wrd is the essential regulatory subunit of the PP2A holoenzyme that antagonises Crb-mediated Ex phosphorylation and degradation.

Since the modulation of Ex^1-468^ protein stability in conditions where ectopic Crb is absent appears to have additional layers of regulation, we also characterised the PP2A holoenzyme(s) that mediate this process. RNAi-mediated depletion of *wrd* suppressed the stabilisation of Ex protein levels induced by Mts expression (Mts^BL^, Fig. 4A-C,I) in the posterior compartment of the wing disc. Expression of Mts^R268A^, the Mts mutant that affects binding to Wrd, did not result in increased levels of the Ex reporter, which was indistinguishable from controls (Fig. 4A,F,G,J), therefore reiterating the role of Wrd in Crb-mediated regulation of Ex protein stability. Additionally, expression of tws-RNAi also suppressed Mts^BL^-mediated stabilisation of Ex (Fig. 4A,B,E,I), thereby suggesting that Tws is also involved in maintaining Ex protein levels.

Furthermore, much like Mts^WT^, expression of Mts^L186A^ in the posterior compartment resulted in an increase in Ex reporter levels (Fig. 4A,F,H,J). This suggests that Cka and, by extension the STRIPAK complex, are not part of the PP2A holoenzyme that regulates Ex stability in conditions where Crb is absent. Together, these results demonstrate that although only Wrd is essential for PP2A-mediated stabilisation of Ex in the presence of Crb, both Wrd and Tws are involved in Crb-independent regulation of Ex proteostasis.

Next, we wanted to explore whether Mts and the identified PP2A holoenzymes physically interact with Ex and if this interaction is modulated by Crb (Fig. 4K, Fig. S6). Using co-IP analyses, we observed that Mts interacts with Ex, both in the presence and absence of Crb^intra^. Interestingly, in the absence of Crb^intra^, Ex primarily interacted with Tws. However, in the presence of Crb^intra^, Ex bound Tws and Wrd. The Ex:Tws interaction observed in the absence of Crb is consistent with the role played by Tws in regulating Ex protein levels *in vitro* and *in vivo*. Additionally, since ***tws*** depletion did not affect the ability of PP2A to stabilise Ex in presence of Crb^intra^, the Ex:Tws interaction detected in the presence of Crb^intra^ is likely to correspond to a potentially constitutively formed complex that is likely unaffected by Crb expression. Furthermore, Wrd associated with Ex only in the presence of Crb^intra^, indicating that the Crb-independent regulation of Ex levels by PP2A^Wrd^ might involve an indirect mechanism, in contrast to what was observed in the context of Crb^intra^. Moreover, these results suggest that, although Crb^intra^ initiates the process leading to Ex phosphorylation and degradation, it simultaneously sets in motion a molecular mechanism that limits the extent to which Ex is degraded, by promoting the Ex:PP2A^Wrd^ interaction that blocks depletion of Ex protein levels.

### Mer and Kib regulate Ex protein levels in a post-translational manner

Mer and Kib have been shown to interact and form complexes with Ex at the apical junction to activate the Hpo kinase cascade (Genevet *et al*., 2010; Yu *et al*., 2010; Genevet and Tapon, 2011; Sun, Reddy and Irvine, 2015; Tokamov *et al*., 2021). More recently, it was shown that Mer and Kib spatially localise to the medial cortex and can activate the Hpo pathway independently of Ex (Su *et al*., 2017). Thus, we were interested to determine the role played by these components in regulating Ex levels post-translationally. We observed that expression of Mer in S2 cells stabilised Ex^1-468^ both in the presence and absence of the Crb stimulus (Fig. 5A). However, Kib was only able to stabilise Ex in the absence of Crb (Fig. 5A). Remarkably, we found that Kib associated with Mts (Fig. 5C), whilst no interaction was observed between Mts and Mer (Fig. 5B). Since Kib was only involved in the regulation of Ex steady-state levels and appeared to interact with PP2A, we next tested whether Kib could also interact with Wrd and Tws, the regulatory subunits that regulate Ex levels in the absence of Crb function (Fig. 5D). Interestingly, we observed that Kib specifically bound to Wrd, but not Tws, thereby suggesting that Kib may regulate Ex levels via PP2A^Wrd^. This result also reinforces the idea that although both PP2A^Wrd^ and PP2A^Tws^ regulate Ex protein levels in a Crb-independent manner, the underlying mechanisms involved in this regulation may be distinct.

**Fig 5.**
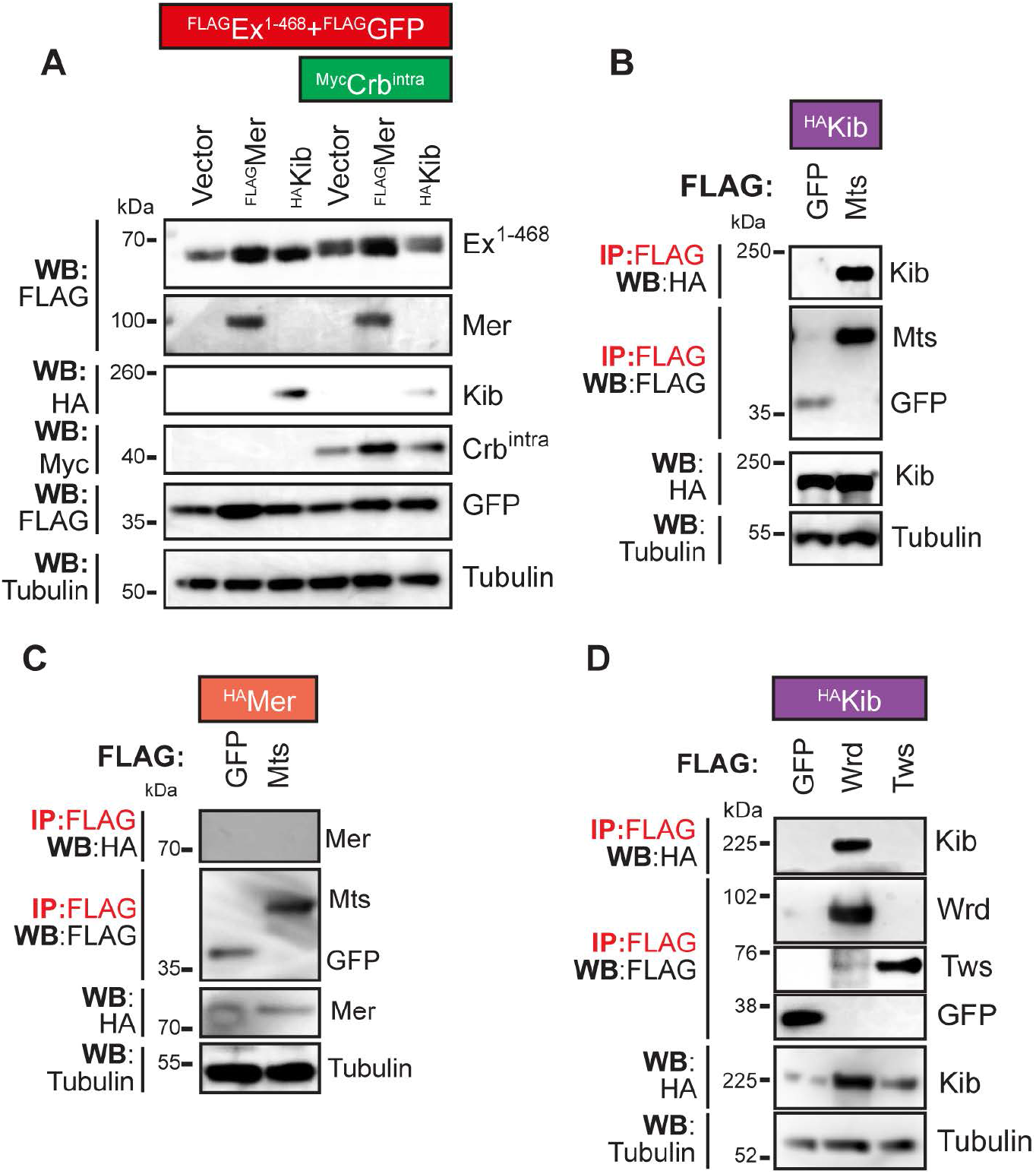
Kib interacts with PP2A ^Tws^ and regulates Ex. **(A)** Mer- and Kib-mediated regulation of Ex protein stability. S2 cells were transfected with the indicated constructs 48h prior to cell lysis and immunoblot analysis with the indicated antibodies. Kib and Mer stabilise Ex in the absence of Crb^intra^. Mer, but not Kib, stabilises Ex in the presence of Crb^intra^. **(B-D)** S2 cells were transfected with the indicated constructs. Following lysis, FLAG-tagged proteins were purified using FLAG-agarose beads. Lysates and FLAG immunoprecipitates were processed for Western blotting analysis with the indicated antibodies. (B) Mts co-immunoprecipitates with Kib. (C) Mts does not interact with Mer. (D) Wrd, but not Tws, binds to Kib.

**Fig 6.**
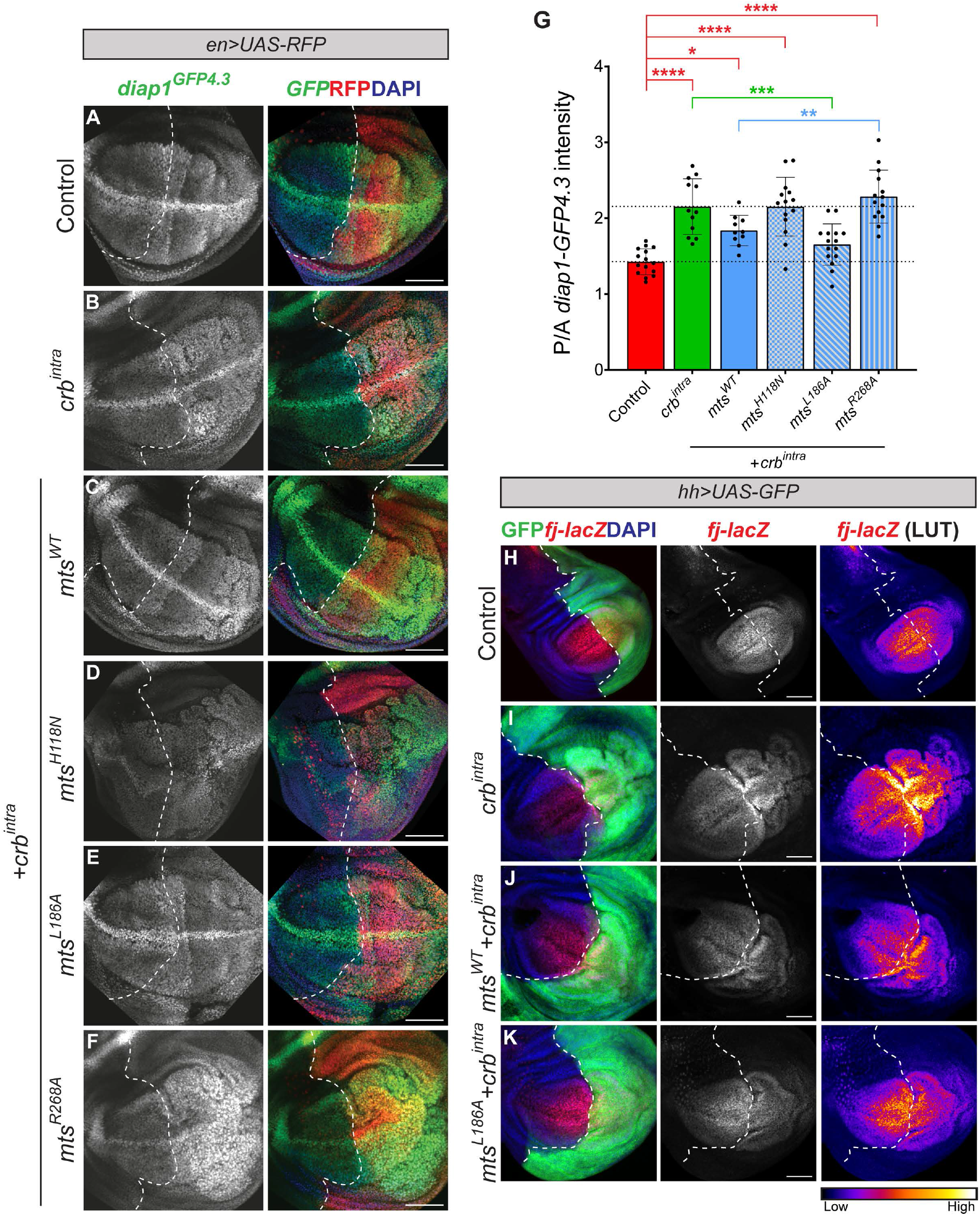
PP2A ^Wrd^ holoenzyme inhibits the Hpo pathway inactivating effects of Crb ^intra^. **(A-F)** PP2A-mediated regulation of Hippo signalling *in vivo* readout, *diap1*^*GFP4*.*3*^. Confocal micrographs show XY sections of third instar wing imaginal discs expressing *diap1*^*GFP4*.*3*^ (green) and UAS-RFP (red) alone (control, A) or in combination with indicated transgenes in the posterior compartment, under the control of the *en-Gal4* driver. Nuclei were stained with DAPI (blue). Dashed white lines depict the AP boundary. Scale bars correspond to 50μm. **(G)** Quantification of relative *diap1*^*GFP4*.*3*^*in vivo* reporter levels. Bar chart depicts the mean ratio of posterior to anterior *diap1*^*GFP4*.*3*^intensity for the indicated genotypes, with all data points represented. Significance was assessed by a one-way ANOVA. Sidak’s multiple comparisons post-hoc test were conducted comparing control, *crb* ^*intra*^ and *mts* ^*WT*^ +*crb* ^*intra*^ conditions against all other genotypes and only significant comparisons are shown. n≥10 for all genotypes. *, p<0.05; **, p<0.01; ***, p<0.001; ****p<0.0001. Note that overexpression of *crb* ^*intra*^ results in increased *diap1*^*GFP4*.*3*^levels. Expression of *mts* ^*L186A*^ abrogates the *crb* ^*intra*^-induced increase, whereas *mts* ^*R268A*^ has no effect. **(H-K)** PP2A-mediated regulation of *fj-lacZ in vivo* Hippo pathway activity reporter. Confocal micrographs show XY sections of third instar wing imaginal discs expressing GFP (green) and *fj-lacZ* (red) alone (control, H) or in combination with indicated transgenes in the posterior compartment, under the control of the *hh-Gal4* driver. Due to the gradient expression of *fj-lacZ* (high in wing pouch and low at the hinge), Fire LUT was applied to the corresponding *fj-lacZ* channel (right). Nuclei were stained with DAPI (blue). Dashed white lines depict the AP boundary. Scale bars correspond to 50μm.

### PP2A ^Wrd^ activates Hpo signalling in the context of Crb expression

Overexpression of Crb^intra^ has been associated with an increase in Yki activity reporter levels such as ex-lacZ and *diap1-GFP* in wing discs (Sherrard and Fehon, 2015; Zhang *et al*., 2015), which is consistent with its role in inhibiting the Hpo pathway. Additionally, knocking down components of the STRIPAK complex, including the Cka regulatory subunit, resulted in a decrease in *ex-lacZ*, since the STRIPAK complex inhibits the Hpo pathway (Ribeiro *et al*., 2010). Having determined that PP2A^Wrd^ antagonises the effects of Crb^intra^ on Ex protein levels and given that PP2A^Cka^ is involved in inhibiting Hpo phosphorylation, we wanted to assess the effects of this Ex-stabilising PP2A complex on Hpo signalling activity.

We monitored Hpo pathway activity *in vivo* using a Yki-responsive *diap1* ^*GFP4*.*3*^ reporter, diap1. As expected, we observed that overexpression of *UAS-crb* ^*intra*^ in the posterior compartment of the wing disc (under the control of *en-GAL4*), resulted in an increase in *diap1* ^*GFP4*.*3*^ levels (Fig. 6A,B,G). When *mts* ^*WT*^ and *crb* ^*intra*^ were co-expressed in the posterior compartment, *diap1* ^*GFP4*.*3*^ levels were elevated compared to controls, but not to the extent observed with *crb* ^*intra*^ alone. Although the mean *diap1* ^*GFP4*.*3*^ levels of *mts* ^*WT*^ +*crb* ^*intra*^ were lower than those of *crb* ^*intra*^, this difference was not statistically significant when all genotypes were included in the comparisons, but only when the Control, *crb* ^*intra*^ and *mts* ^*WT*^ +*crb* ^*intra*^ conditions were considered. Taken together, this suggests that Mts^WT^ may partially abrogate the increase in Yki activity induced by Crb^intra^ (Fig. 6A-C,G). In contrast, co-expression of *mts*^*H118N*^ and *crb*^*intra*^ did not affect the ability of Crb^intra^ to increase Yki activity, demonstrating the specificity of the effects observed with overexpression of Mts^WT^ and Crb^intra^ (Fig. 6A,B,D,G). We also tested the effects of PP2A on another Yki reporter, *four-jointed* (*fj*)-lacZ. We observed that overexpression of Crb^intra^ in the posterior compartment under the control of *hh-Gal4*, resulted in an increase in *fj-lacZ* levels (Fig. 6H,I). Furthermore, when *mts* ^*WT*^ and *crb* ^*intra*^ were co-expressed, we observed lower *fj-lacZ* levels in the posterior compartment when compared with expression of *crb* ^*intra*^ alone (Fig. 6I,J), similar to results observed with the *diap1*^*GFP4*.*3*^ reporter.

To decouple the effects of PP2A^Wrd^ and PP2A^Cka^ on Hpo pathway activity, we used the Mts^L186A^ (which prevents PP2A^Cka^ holoenzyme assembly and therefore would promote higher levels of the PP2A^Wrd^ complex) and Mts^R268A^ (which abrogates PP2A^Wrd^ holoenzyme assembly and would be predicted to lead to increased PP2A^Cka^ levels) mutants. When *mts*^*L186A*^ was co-expressed with *crb* ^*intra*^, we observed that this completely abrogated the increase in *diap1* ^*GFP4*.*3*^ levels induced by expression of only *crb* ^*intra*^ (Fig. 6A,B,E,G). These findings indicate that in the absence of the STRIPAK complex, PP2A likely containing the Wrd subunit, can inhibit the Crb-dependent increase in Yki activity. Additionally, co-overexpression of *mts* ^*L186A*^ and *crb* ^*intra*^, resulted in reduced intensity of *fj-lacZ* in the posterior compartment (Fig. 6K), mirroring the effects observed with the *diap1*^*GFP4*.*3*^ reporter. Remarkably, co-expression of *mts*^*R268A*^ and *crb*^*intra*^ did not affect the ability of Crb^intra^ to increase *diap1*^*GFP4*.*3*^ levels (Fig. 6A,B,F,G). This suggests that Wrd is essential for Mts to be able to antagonise the effect of Crb on Yki activity. Taken together, these results demonstrate that PP2A^Wrd^, the holoenzyme that stabilises Ex, abrogates the increase in Yki activity induced by Crb^intra^, by activating the Hpo pathway.

## Discussion

Given its crucial role in the Hpo pathway, Ex protein levels are tightly regulated by multiple post-translational mechanisms that modulate its localisation and stability (Ling *et al*., 2010; Robinson *et al*., 2010; Chen *et al*., 2011; Ribeiro *et al*., 2014; Zhang *et al*., 2015; Su *et al*., 2017; Ma *et al*., 2018; Fulford *et al*., 2019; Song and Ma, 2023). The apicobasal polarity protein Crb precisely regulates Ex function by both facilitating its recruitment to the apical membrane and limiting Ex activity by promoting its phosphorylation and subsequent ubiquitin-mediated degradation (Ribeiro *et al*., 2014; Fulford *et al*., 2019). However, the mechanisms that ensure this balance is precisely maintained remain largely unknown. Here, we show that the PP2A^Wrd^ holoenzyme dephosphorylates and stabilises Ex in the presence of Crb. This role of PP2A occurs independently of Cka and, thus, of the STRIPAK complex. Furthermore, in contrast to the previously reported inhibitory role of PP2A^Cka^, we demonstrate that PP2A^Wrd^ activates the Hpo pathway, limiting Yki activity induced by Crb. These results elucidate a critical PP2A nexus that maintains Ex levels in a delicate equilibrium to precisely regulate Hpo signalling activity (Fig. 7).

**Fig 7.**
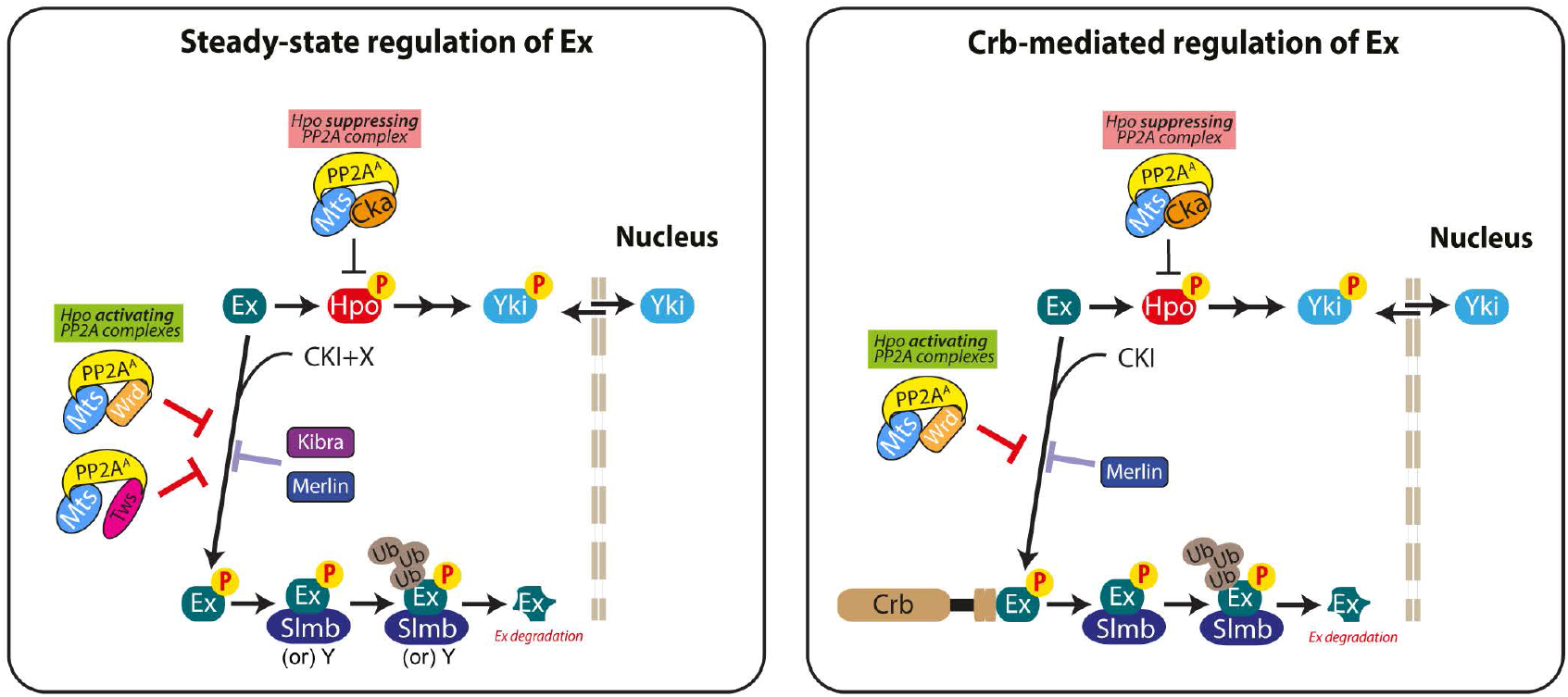
Proposed model for the dual role of PP2A in the regulation of Ex protein levels. Schematic detailing the key findings from this study and proposed model. In steady-state, CKI and unknown kinase(s), X, can promote Ex phosphorylation, which subsequently results in degradation of Ex in a manner partly dependent on Slmb; Y represents an uncharacterised mechanism that exists in regulating homeostatic Ex. PP2A^Wrd^, PP2A^Tws^, Mer and Kib are involved in maintaining steady-state Ex levels. In the presence of the Crb stimulus, PP2A^Wrd^, antagonises Crb-dependent phosphorylation and degradation of Ex, and hence, activates the Hpo pathway.

We observed that, in steady-state, the CKI kinases Ck1α and Gish promote phosphorylation of Ex, and this can be reversed by PP2A. Moreover, Slmb is also involved in Ex degradation in these conditions. Interestingly, in the absence of ectopic Crb, we observed that the regulation of Ex by Ck1α, Gish and Slmb is limited to the Slmb consensus site ^452^TSGIVS^457^, which is similar to the previously characterised mechanism involved in Crb-dependent regulation of Ex (Ribeiro *et al*., 2014; Fulford *et al*., 2019). Remarkably, an Ex truncation lacking the ^452^TSGIVS^457^ site can be stabilised by PP2A despite being refractory to CKI kinase-dependent phosphorylation and Slmb-dependent degradation, thereby suggesting the involvement of other kinases and ubiquitin ligases in the regulation of Ex stability, at least in the absence of Crb. Furthermore, we observed that PP2A can promote stability of Ex^FL^ and mapped this PP2A-mediated regulation of Ex to the N-terminus of Ex (Ex^1-468^). Although Ck1α and Gish phosphorylate Ex both in the context of a Crb-dependent and -independent mechanism, whether Ex is phosphorylated to a higher extent when associated with Crb is not currently known. CKIs prefer substrates that are primed by prior phosphorylation events (Cheong and Virshup, 2011; Cruciat, 2014), hence the phosphorylation events involved in the regulation of Ex proteostasis in the absence of Crb could prime CKI-mediated phosphorylation of Ex to rapidly adapt to dynamic polarity cues.

We showed that the PP2A^Wrd^ and PP2A^Tws^ holoenzymes are essential for maintaining Ex steady-state protein levels. Since PP2A regulatory subunits typically recruit and dephosphorylate different substrates, the involvement of both regulatory subunits in PP2A-mediated stabilisation of Ex in conditions where Crb function is dispensable was unexpected. This observation parallels findings where both mammalian B56 (Wrd) and B55 (Tws) regulatory subunit families have been implicated in dephosphorylation of common targets, including Tau (Yu, Yoo and Ahn, 2014), the NF-κB pathway inhibitory kinase IκBα (Tsuchiya *et al*., 2017) and the protein kinase Akt (Rodgers, Vogel and Puigserver, 2011; Ruvolo *et al*., 2011). In these cases, the B55 and B56 regulatory subunits employ distinct mechanisms to regulate the common substrates. Interestingly, we observed that in conditions where no ectopic Crb is present in cells, Ex binds only to Tws, suggesting that Wrd and Tws employ specific molecular mechanisms to regulate Ex.

In the presence of Crb, we observed a shift in the association of Ex from Tws to Wrd, which complements our observation that PPA^Wrd^ functionally antagonises Crb-mediated Ex degradation. The differential binding of Wrd to Ex observed in the absence and presence of Crb, suggests that the PP2A^Wrd^ holoenzyme may stabilise Ex through different mechanisms in these conditions, even though a similar overall effect on Ex stability is achieved. For instance, in the absence of the Crb stimulus, PP2A^Wrd^ may regulate an unknown upstream kinase or a negative regulator of Ex, thereby influencing Ex stability without directly associating with it. These results demonstrate that multiple PP2A complexes regulate Ex stability in a context-dependent manner.

Kib, Ex and Mer have been shown to form a complex that acts upstream of the core Hpo components to activate the Hpo pathway (Boggiano and Fehon, 2012; Enderle and McNeill, 2013). More recently, Kib and Mer have been shown to activate the Hpo pathway in a spatially distinct manner from Ex. Kib and Mer recruit Hpo and Wts to the non-junctional medial cortex, while Ex localises with Crb to the junctional cortex (Su *et al*., 2017). Additionally, Crb represses the Hpo- and growth-regulating functions of Kib by sequestering Kib to the junctions, away from the medial apical cortex (Su *et al*., 2017). Interestingly, we observed that Mer regulated Ex protein levels both in the absence and in the presence of the Crb stimulus. However, Kib only regulated Ex protein levels when Crb was not present, which is in accordance with the role of Crb in repressing Kib function. Hence, these results reinforce the importance of the relationship between Mer, Kib and Ex. Although previous reports stated that Kib does not affect Ex levels *in vivo* (Su *et al*., 2017), these experiments relied on Ex antibody staining rather than assessing post-translational effects using the *ubi-Ex1*^*-468*^*::GFP* stability reporter. Therefore, it would be interesting to determine the role of Kib in the regulation of Ex protein stability *in vivo*, using the approaches described in the current study. Remarkably, we observed that Kib binds to Mts and Wrd and, to the best of our knowledge, this is the first study to report an interaction between PP2A and Kib. Therefore, in conditions where Crb function is absent, Kib could regulate Ex stability by mediating the function of the PP2A^Wrd^ holoenzyme to stabilise Ex. Crb would disrupt the formation of this complex, thereby preventing the stabilisation of Ex by the Kib-PP2A^Wrd^ complex. This model suggests that Kib recruits PP2A^Wrd^ to phosphorylated Ex to promote PP2A-mediated dephosphorylation and stabilisation of Ex. However, in the presence of Crb, this function of Kib would be repressed, thereby resulting in the phosphorylation and degradation of Ex.

Cell polarity is an essential determinant of epithelial architecture, and the Hpo pathway plays a key role in integrating polarity cues to mediate tissue growth (Campbell, Knust and Skaer, 2009; Sherrard and Fehon, 2015; Campanale, Sun and Montell, 2017; Peglion and Goehring, 2019). Our work shows that distinct PP2A complexes intricately regulate Ex protein levels in steady-state conditions and in response to changes in Crb function. We propose that this context-dependent regulation of Ex is crucial for cells to quickly respond to dynamic polarity cues during tissue development and remodelling.

## Acknowledgements

We thank the Vienna Drosophila Resource Center for providing transgenic RNAi fly stocks used in this study. Stocks obtained from the Bloomington *Drosophila* Stock Center (NIH P40OD018537) were used in this study. The antibodies E7 and 2A1 were deposited by M. Klymkowsky and R. Holmgren, respectively to the Developmental Studies Hybridoma Bank, created by the NICHD of the NIH and maintained at The University of Iowa, Department of Biology. We thank N. Tapon for providing fly stocks. We thank Sam Wallis and Linda Hammond from the Barts Cancer Institute Microscopy Service for assistance with microscopy. We thank Olivia Wakefield and Guanxiang Hua for assistance in preliminary experiments. We thank members of the Ribeiro lab for helpful discussions and N. Tapon, M. Holder, J. Marshall and S. Martin for critical reading of the manuscript. The authors declare no conflicts of interest. This work was supported by funding from Cancer Research UK (C16420/A18066), The Academy of Medical Sciences/Wellcome Trust Springboard Award (SBF001/1018), The Brain Tumour Charity (GN-000408) and from the Biotechnology and Biological Sciences Research Council (BB/T004576/1). A.R. was supported by a CRUK PhD studentship (S_3967).

## Figure Legends

**Fig S1. List of PP2A mutants generated in this study**

Protein sequence alignment of Mts (Dm_mts) and human PP2Acα (Hs_PP2ACalpha) with mutations used in this study marked in red. Table lists Mts mutants generated in this study, their predicted effect and their true effect validated in *Drosophila* S2 cells.

**Fig S2. PP2A-mediated regulation of Arm and of Ex in the presence of Crb** ^**intra**^

**(A)** Assessment of PP2A catalytic activity *in vitro*. S2 cells were transfected with the indicated constructs 48h prior to cell lysis and processing for immunoblot analysis with the indicated antibodies. Note that Arm is constitutively degraded in S2 cells in a phosphorylation-dependent manner. Mts^WT^ and Mts^R268A^ stabilise Arm^WT^, therefore indicating that Mts^R268A^ is catalytically active. Mts^H118N^ is unable to stabilise Arm^WT^, demonstrating that Mts^H118N^ is catalytically inactive. Arm^S>A^ contains a mutation in the Arm phosphorylation site rendering it refractory to protein degradation. **(B)** PP2A stabilises Ex in the presence of Crb^intra^ in a dose-dependent manner. GFP and Tubulin were used as transfection and loading controls, respectively. **(C-E)** Mts-mediated regulation of Ex *in vivo* reporter. Confocal micrographs show XY (top) and transverse sections (bottom) of third instar wing imaginal discs expressing *ubi-Ex*^*1-468*^ *::GFP* (green) and *UAS-mIFP* (red) alone (control, C) or in combination with indicated transgenes in the posterior compartment, under the control of the *hh-Gal4* driver. Nuclei were stained with DAPI (blue). Dashed white lines depict the AP boundary. Scale bars correspond to 50μm. **(F)** Quantification of Ex *in vivo* reporter and effect of Mts^BL^ expression. Bar chart depicts the geometric mean ratio of posterior to anterior Ex^1-468^::GFP intensity for the indicated genotypes, with all data points represented. Significance was assessed by a one-way ANOVA conducted on log-transformed data, followed by Tukey’s multiple comparisons post-hoc test. n≥17 for all genotypes. ****, p<0.0001. Expression of mts^BL^ rescued Ex protein levels from Crb-dependent degradation.

**Fig S3. Assessing the role of Cka in Crb-dependent regulation of Ex**

**(A-D)** Role of Cka in Ex regulation. Confocal micrographs show XY (top) and transverse sections (bottom) of third instar wing imaginal discs expressing *ubi-Ex*^*1-468*^ *::GFP* (green) and UAS-mIFP (red) alone (control, A) or in combination with indicated transgenes in the posterior compartment, under the control of the *hh-Gal4* driver. Crosses were reared at 18°C. Nuclei were stained with DAPI (blue). Dashed white lines depict the AP boundary. Scale bars correspond to 50μm. **(E)** Quantification of Ex reporter levels. Bar chart depicts the mean ratio of posterior to anterior Ex^1-468^::GFP intensity for the indicated genotypes, with all data points included. Significance was assessed by a one-way ANOVA. n≥10 for all genotypes. **(F)** Cka interaction with PP2A catalytic subunit *in vitro*. S2 cells were transfected with the indicated constructs. Following lysis, FLAG-tagged proteins were purified using FLAG-agarose beads. Lysates and eluates were processed for Western blotting analysis using the indicated antibodies. **(G)** Quantification of Cka:Mts interaction. Bar chart depicts the levels of Cka bound to Mts^L186A^ relative to Mts^WT^ per experiment. Significance was assessed by a paired t test. n=3 independent experiments. **, p<0.01. **(H)** Assessment of catalytic activity of STRIPAK-deficient PP2A mutant. S2 cells were transfected with the indicated constructs 48h prior to cell lysis and processing for immunoblot analysis with the indicated antibodies. Mts^L186A^ stabilises Arm^WT^, indicating that Mts^L186A^ is catalytically active.

**Fig S4. PP2A regulates Ex proteostasis via its N-terminal region**

If mentioned, S2 cells were treated with indicated dsRNA for 24h prior to transfection with the indicated constructs. (A) Knocking down mts reduces steady-stateEx^1-468^ protein levels. (B) In the absence of Crb, Mts was able to increase Ex levels, while none of the catalytic subunits of the other phosphatases influenced steady-state Ex levels. (C) Mts^WT^ did not have any effect on Ex^469-1030^ and Ex^1031-1427^ levels. (D) Mts^WT^, but not Mts^H118N^, was able to increase Ex^FL^ protein levels in conditions where Crb function is dispensable.

**Fig S5. Validation of dsRNAs and Mts** ^**R268A**^

**(A)** Validation of *wdb* and *wrd* dsRNA efficiency. S2 cells were treated with the indicated dsRNA for 24h prior to transfection with the indicated constructs. Cells were lysed 48h after transfection and lysates were processed and analysed by Western blotting using the indicated antibodies. GFP and Tubulin were used as transfection and loading controls, respectively. *wrd* and *wdb* dsRNAs specifically downregulate *Wrd* and *Wdb* protein levels. **(B)** Validation of *tws* dsRNA reagents. Since *tws* dsRNA targets the 3’UTR of the *tws* gene, RT-PCR was conducted on lysates from S2 cells treated with either lacZ dsRNA or *tws* dsRNA. RT-PCR reactions were run on a 2% agarose gel. *tws* dsRNA resulted in a reduction in *tws* RNA levels. rp49 was used as loading control. **(C-E)** Effect of R286A mutation on the Mts:Wrd and Mts:Wdb interactions. S2 cells were transfected with the indicated constructs. Following lysis, FLAG-tagged proteins were purified using FLAG-agarose beads. Lysates and eluates were processed for Western blot analysis with the indicated antibodies. Tubulin was used as loading control. (D,E) Bar charts depicting the levels of either Wrd (D) or Wdb (E) bound to Mts^R268A^ relative to Mts^WT^ per experiment. Significance was assessed by a paired t test and only significant comparisons were shown. n=3 independent experiments. *, p<0.05.

**Fig S6. Mts binds Ex in the absence or presence of Crb expression**

**(A)** Effect of Crb on the Mts:Ex interaction. S2 cells were transfected with the indicated constructs and lysed 48h after transfection. FLAG-tagged proteins were purified from lysates using FLAG-agarose beads. Lysates and FLAG immunoprecipitates were processed for immunoblot analysis using the indicated antibodies. Tubulin was used as loading control. Mts co-immunoprecipitated with Ex^1-468^ in the presence and absence of Crb^intra^.

