## Supplementary Figures for "A Dual Role for the PP2A Phosphatase in Hippo Signalling Regulation"

### Figure S1

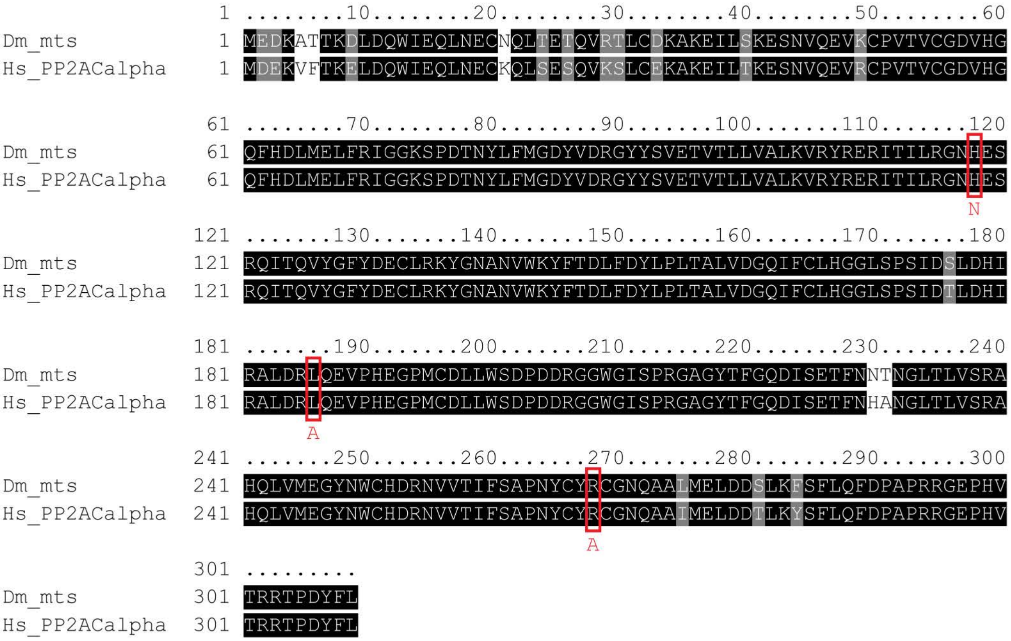


| Mts Mutation | Predicted Effect | Validated Effect |
|---|---|---|
| H118N | Loss of catalytic activity | Loss of catalytic activity (by Arm stabilisation assay) |
| R268A | Block interaction with Wrd/Wdb | Block interaction with Wrd (by co-IP) |
| L186A | Block interaction with Cka | Block interaction with Cka (by co-IP) |

**Figure S2**

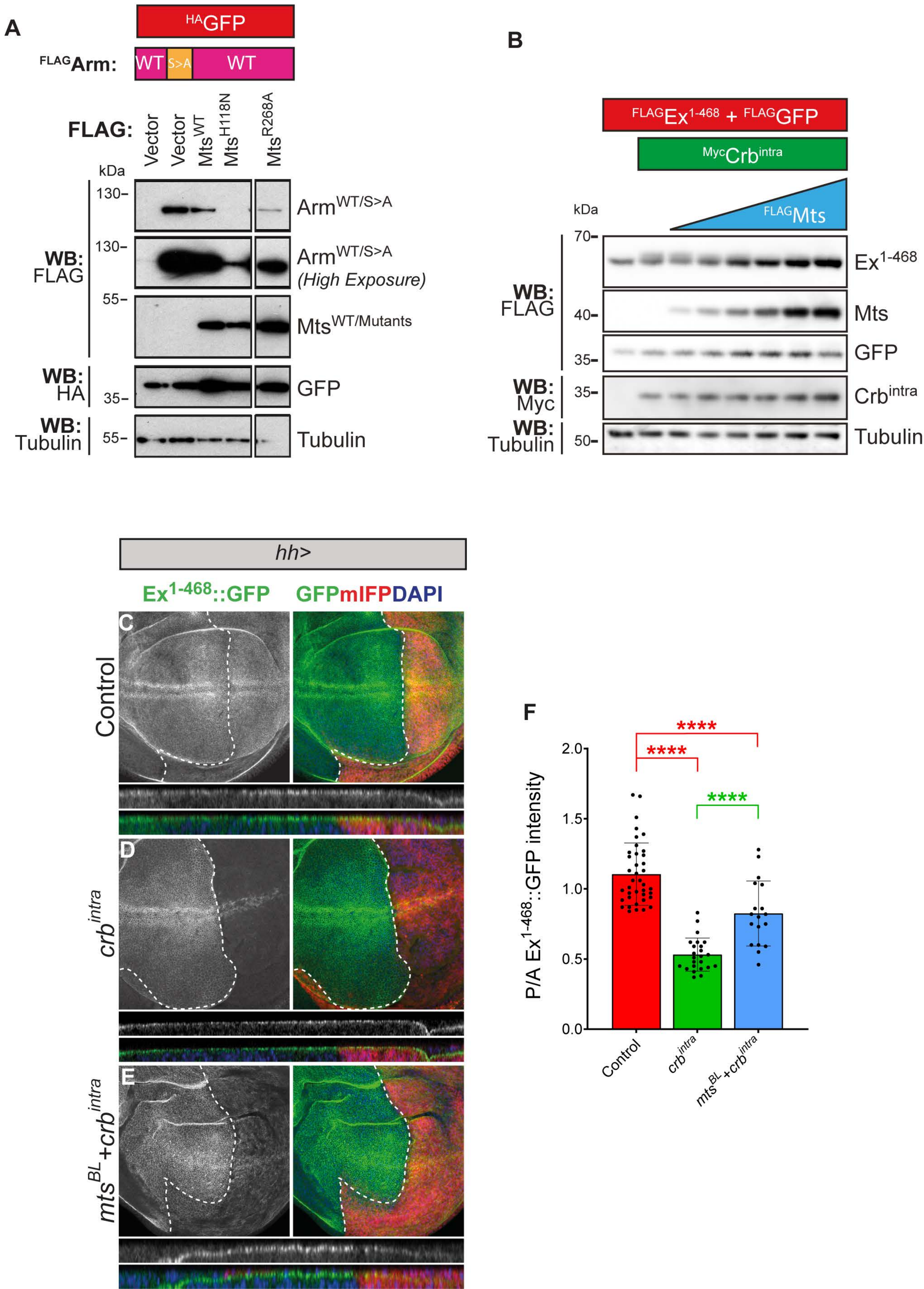
A
HAGFP
FLAGArm: WT S>A WT
FLAG: Vector Vector MtsWT MtsH118N MtsR268A
kDa
130
130
55
35
55
WB: FLAG
WB: HA
WB: Tubulin
ArmWT/S>A
ArmWT/S>A (High Exposure)
MtsWT/Mutants
GFP
Tubulin
B
FLAGEx1-468 + FLAGGFP
MycCrbintra
FLAGMts
kDa
70
40
35
35
50
WB: FLAG
WB: Myc
WB: Tubulin
Ex1-468
Mts
GFP
Crbintra
Tubulin
hh>
Ex1-468::GFP
GFPmIFPDAPI
C
D
E
Control
crbintra
mtsBL+crbintra
F
****
****
****
P/A Ex1-468::GFP intensity
2.0
1.5
1.0
0.5
0.0
Control
crbintra
mtsBL+crbintra

**Figure S3**

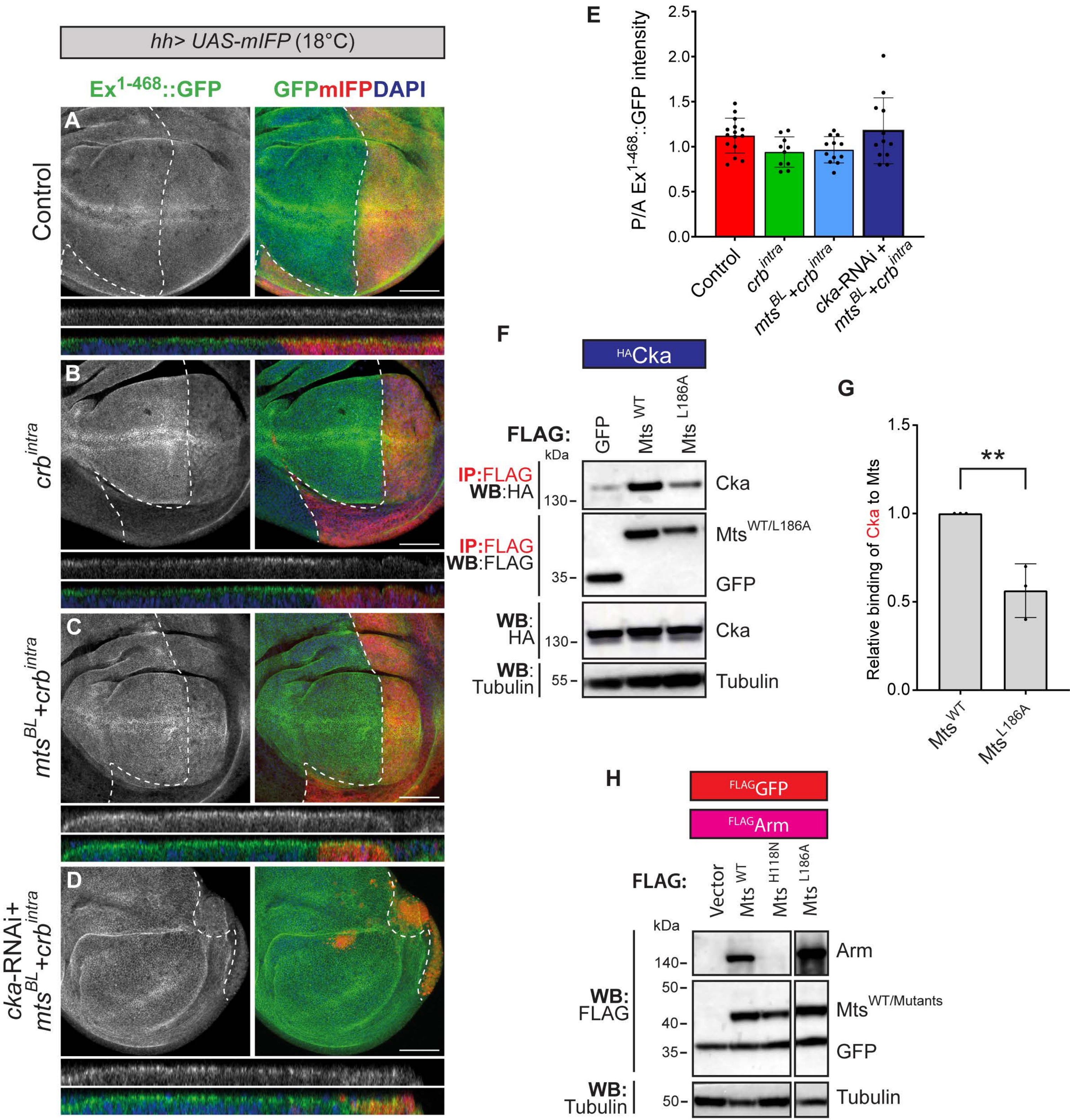

hh> UAS-mIFP (18°C)
Ex1-468::GFP
GFPmIFPDAPI
A
B
C
D
Control
crb intra
mts BL +crb intra
cka-RNAi+ mts BL +crb intra
E
P/A Ex1-468::GFP intensity
2.5
2.0
1.5
1.0
0.5
0.0
Control
crb intra
mts BL +crb intra
cka-RNAi + mts BL +crb intra
F
HACka
FLAG:
GFP
Mts WT
Mts L186A
kDa
IP:FLAG WB:HA
130
Cka
IP:FLAG WB:FLAG
35
Mts WT/L186A
GFP
WB: HA
130
Cka
WB: Tubulin
55
Tubulin
G
Relative binding of Cka to Mts
1.5
1.0
0.5
0.0
**
Mts WT
Mts L186A
H
FLAGGFP
FLAGArm
FLAG:
Vector
Mts WT
Mts H118N
Mts L186A
kDa
140
Arm
WB: FLAG
50
40
35
Mts WT/Mutants
GFP
WB: Tubulin
50
Tubulin

### Figure S4

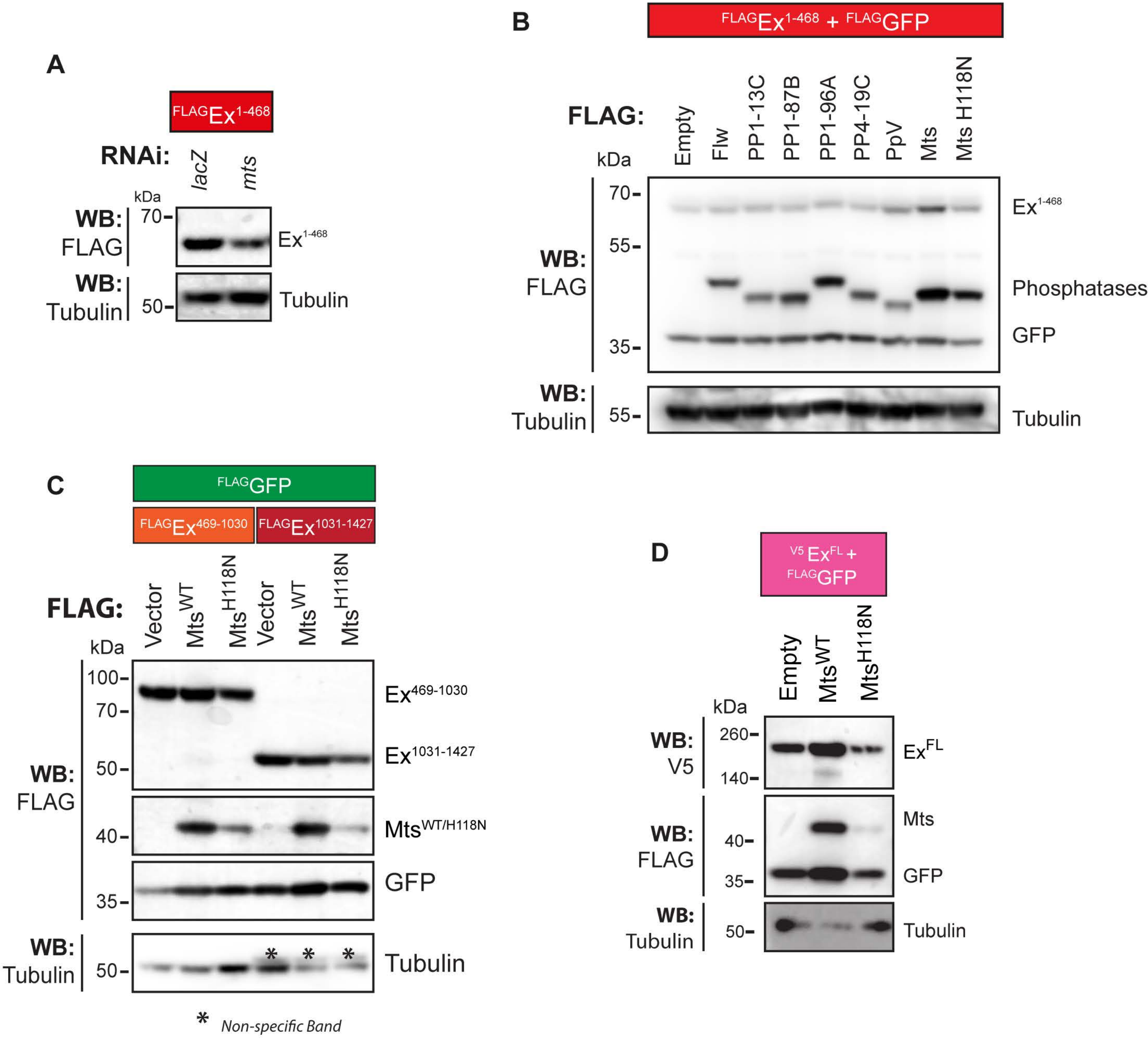
A
FLAGEx1-468
RNAi:
lacZ
mts
kDa
WB: FLAG
70
Ex1-468
WB: Tubulin
50
Tubulin
B
FLAGEx1-468 + FLAGGFP
FLAG:
Empty
Flw
PP1-13C
PP1-87B
PP1-96A
PP4-19C
PpV
Mts
Mts H118N
kDa
70
55
35
WB: FLAG
Ex1-468
Phosphatases
GFP
WB: Tubulin
55
Tubulin
C
FLAGGFP
FLAGEx469-1030
FLAGEx1031-1427
FLAG:
Vector
MtsWT
MtsH118N
Vector
MtsWT
MtsH118N
kDa
100
70
50
40
35
WB: FLAG
Ex469-1030
Ex1031-1427
MtsWT/H118N
GFP
WB: Tubulin
50
Tubulin
* Non-specific Band
D
V5 ExFL + FLAGGFP
Empty
MtsWT
MtsH118N
kDa
260
140
WB: V5
ExFL
WB: FLAG
40
35
Mts
GFP
WB: Tubulin
50
Tubulin

Figure S5

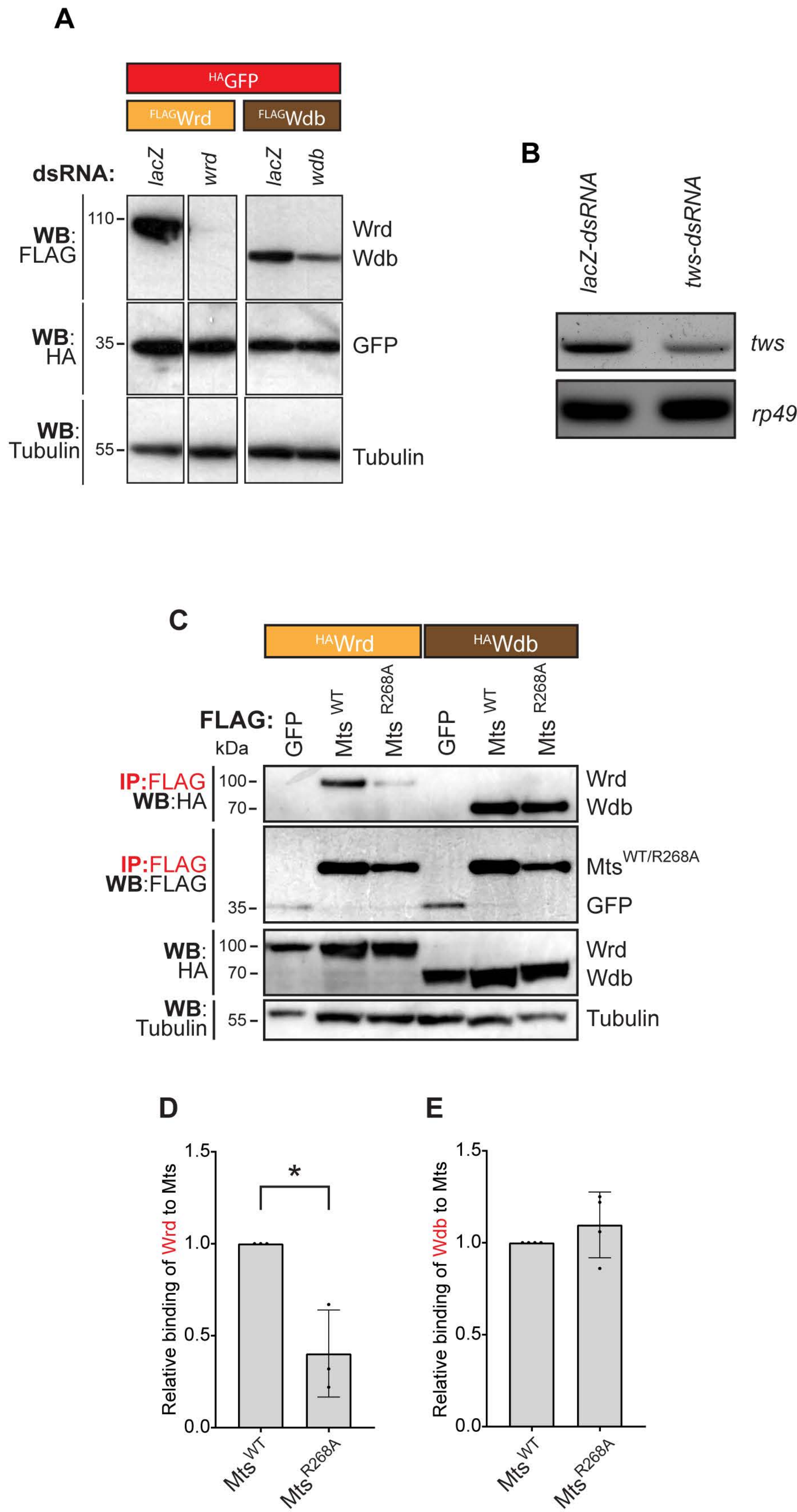
A
HAGFP
FLAGWrd
FLAGWdb
dsRNA:
lacZ
wrd
lacZ
wdb
WB: FLAG
110
Wrd
Wdb
WB: HA
35
GFP
WB: Tubulin
55
Tubulin
B
lacZ-dsRNA
tws-dsRNA
tws
rp49
C
HAWrd
HAWdb
FLAG:
GFP
Mts WT
Mts R268A
GFP
Mts WT
Mts R268A
kDa
IP:FLAG
WB:HA
100
70
Wrd
Wdb
IP:FLAG
WB:FLAG
Mts WT/R268A
35
GFP
WB: HA
100
70
Wrd
Wdb
WB: Tubulin
55
Tubulin
D
Relative binding of Wrd to Mts
1.5
1.0
0.5
0.0
*
Mts WT
Mts R268A
E
Relative binding of Wdb to Mts
1.5
1.0
0.5
0.0
Mts WT
Mts R268A
